# Differential impact of plant secondary metabolites on the soil microbiota

**DOI:** 10.1101/2021.02.09.430398

**Authors:** Vadim Schütz, Katharina Frindte, Jiaxin Cui, Pengfan Zhang, Stéphane Hacquard, Paul Schulze-Lefert, Claudia Knief, Margot Schulz, Peter Dörmann

## Abstract

Plant metabolites can shape the microbial community composition in the soil. Two indole metabolites, benzoxazolinone (BOA) and gramine, produced by different Gramineae species, and quercetin, a flavonoid synthesized by many dicot species, were studied for their impacts on the community structure of soil bacteria. The three plant metabolites were directly added to agricultural soil over a period of 28 days. Alterations in bacterial composition were monitored by next generation sequencing of 16S *rRNA* gene PCR products and phospholipid fatty acid analysis. Treatment of the soil with the plant metabolites altered the composition of bacterial taxa on the phylum and genus levels. Alpha diversity was significantly altered by BOA or quercetin, but not by gramine. BOA treatment caused an increase in the relative abundances of only four genera, three of them belonging to the Actinobacteriota. Gramine or quercetin treatment resulted in the increase in relative abundance of 13 or 14 genera, respectively, most of them belonging to the Proteobacteria. The relative abundance of 22 genera was decreased after BOA treatment, 16 of which were also decreased by gramine or quercetin. Isolation and characterization of cultivable bacterial indicated an enrichment in specific *Arthrobacter* or *Pseudomonas* strains. Therefore, the effects of the treatments on soil bacteria were characteristic for each metabolite, with BOA exerting a broad-spectrum inhibitory effect, with only few genera able to proliferate, while gramine and quercetin caused the proliferation of many potentially beneficial strains. As a consequence, benzoxazolinone or gramine biosynthesis which have evolved in different barley species, is accompanied with the association with distinct bacterial communities in the soil, presumably after mutual adaptation during evolution.

## 1 Introduction

Plants establish their specific microbial environment, starting with the attraction of microorganisms from the soil, based on structural carbohydrates and exudates from the roots, after germination, in addition to the seed-born microbiota. Root exudates are crucial for modulating the microbial species composition in the rhizosphere to improve plant growth and the competitiveness with other organisms (Stringlis et al., 2018; Huang et al., 2019; Voges et al., 2019; Harbort et al., 2020). Metabolites and their derivatives exuded from the roots likely contribute to the adjustment of the plant microbiota and modulate its functional diversity. For example, salicylic acid exuded from willow tree roots modulates the microbiota and alters microbial gene expression in the rhizosphere (Schmidt et al., 2000; Lebeis et al., 2015). A considerable amount of allelopathic and microbiota-modifying metabolites is released from rotting plant material, and, in appropriate agricultural culture systems, also from mulches, thereby influencing the soil microbial communities in the entire agricultural ecosystem. These effects have been studied more intensively for selected metabolites, including glucosinolates and their breakdown products (isothiocyanates) (Hollister et al., 2013; Turner et al., 2013; Averill et al., 2014; Hu et al., 2014; Siebers et al., 2018; Hansen et al., 2020), isoprenoids and coumarins (Stringlis et al., 2018; Huang et al., 2019; Voges et al., 2019; Harbort et al., 2020).

Benzoxazinoids (BXs) are a class of indole-derived metabolites with allelochemical activity synthesized in Poaceae, including some *Hordeum* species (e.g. *H. brachyantherum, H. flexuosum, H. lechleri)* (Sicker et al., 2000; Grün et al., 2005). BXs, typically synthesized in young plants, have allelopathic/phytotoxic and other biocidal properties including antinutritional functions in herbivores. BXs are unstable in soil, where they are rapidly converted into benzoxazolinones (BOA, MBOA) which are detectable in the soil for up to 12 weeks. Because of their phytotoxicity, these compounds have been selected for weed control in agriculture. BOA can be degraded or detoxified by a few fungi and bacteria (Schulz et al., 2013; Schulz et al., 2017). Changes in the microbiome structure of the rhizosphere of maize plants have been studied with maize wild type and the BX-deficient *bx1* mutant plants. Cultivation of WT or *bx1* plants resulted in a shift in bacterial and fungal species composition, which affected the growth of the following plant generation (Hu et al., 2018). Several bacterial and few fungal operational taxonomic units (OTUs) in the rhizosphere of maize were influenced by different *bx (bx1, bx2, bx6)* mutations (Cotton et al., 2019). Growth of the same three maize mutants led to a decline in fungal and bacterial species richness in the rhizosphere, with a decrease in many plant pathogens (Kudjordjie et al., 2019). The effects of BOA on microbiota composition and species diversity in the bulk soil remain, however, unclear.

Some Poaceae produce gramine as an additional indole-derived metabolite. Gramine is one of the most important allelochemicals in certain barley cultivars that do not contain BXs *(Hordeum vulgare* L., e.g. genotypes Lina, Osiris) (Larsson et al., 2006; Kokubo et al., 2017). Gramine is also found in other plants, for instance in lupines *(Lupinus luteus* L.). It affects germination and growth of oat, rye and wheat (Bravo et al., 2010). For allelopathic effects, gramine has to be released from rotting plant material, because in contrast to BXs, gramine seems to be absent from root exudates. Gramine also inhibits growth of cyanobacteria and other bacteria (Laue et al., 2014; Popp et al., 2016). The detoxification pathway of gramine in aphids *(Sitobion avenae)* involves carboxylesterase and glutathione S-transferase activities (Cai et al., 2004), but bacterial or plant detoxification strategies are unclear. Barley cultivars produce only one of the two metabolites, BX or gramine, indicative for divergent evolution. Therefore, it is possible that BX or gramine might have distinct effects on the soil microbiota.

Flavonoids are polyphenols which are widespread in plants, in particular in seeds and root exudates, with quercetin representing one of the most abundant flavonoids. Flavonoids are involved in the regulation of nodulation in legumes, they exert antioxidant and antimicrobial activities, and many of them have allelopathic properties. Several flavonoids have a short lifetime while others persist in the soil, depending on the chemical structures, such as the number of functional groups, physicochemical characteristics of the soil and the rate of microbial degradation. Many Fabaceae like *Lotus japonicus* contain quercetin as the main flavonoid (Suzuki et al., 2008). Several microorganisms, such as *Rhizobium, Agrobacterium, Pseudomonas*, *Bacillus* and *Rhodococcus* species degrade flavonoids including quercetin (Pillai and Swarup, 2002). *Rhizobium* species detoxify flavonoids and isoflavonoids via C ring fission, which results in phenolic compounds like protocatechuic acid. The prooxidant activity of protocatechuic acid can cause oxidative stress and death of other microorganisms. Protocatechuic acid is also responsible for the indirect allelopathic effects of catechin, which is e.g. exuded by roots of *Rhododendron formosanum* (Wang et al., 2013). The rhizosphere of *R. formosanum* is rich in microorganisms that use catechin as carbon source, including *Pseudomonas*, *Herbaspirillum* and *Burkholderia* species. These results suggest that flavonoid breakdown products might be the true modifiers of microbial biodiversity and influence the abundance of defined species.

To study the effects of selected plant-root derived metabolites on soil microbial communities in comparison, native agricultural soil was exposed to BOA, gramine or quercetin, using concentrations that occur under natural conditions. We selected BOA and gramine which are two indole metabolites produced in a mutually exclusive manner by Poaceae species, to study their impact on the soil microbiome (Larsson et al., 2006). In addition, we chose to study the effects of quercetin on the soil microbiome with the aim to assess how specific the changes in the soil microbial community introduced by the two indole metabolites are. Bacterial 16S *rRNA* gene-based community profiling in combination with phospholipid fatty acid analyses, revealed that each of the three metabolites exerts a strong and distinct alteration in species diversity and community structure. In addition, microorganisms were retrieved from soil after application of the aforementioned plant specialized compounds with the aim of unraveling whether bacterial genera considered to be tolerant to these compounds after 16S *rRNA* gene community profiling, can be isolated by cultivation.

## 2 Materials Methods

### 2.1 Incubation of soil with secondary metabolites

Freshly harvested, agricultural soil from Cologne (Cologne agricultural soil) (Bai et al., 2015) was sieved (2 mm mesh) and filled into open pots (10 cm diameter) placed onto Petri dishes. An amount of 10 μmol of solid benzoxazolinone (BOA, Fluka Chemika), gramine (Sigma-Aldrich) or quercetin (ABCR, Karlsruhe, Germany) was added and mixed with 300 g of soil (water content of 12% ± 1%). The treatment was repeated every second day over a period of 28 days. In total, 140 μmol of metabolites were added to each pot equivalent to ~0.5 μmol per g soil or ~3.8 mM (based on the water content of 12%). In addition, the soil was watered every second day with germ-free water from the top to maintain a maximal water retention capacity of 70%. Control soil was watered with germ-free water without the addition of metabolites. The pots were covered with transparent plastic covers and incubated in a growth chamber under a 16 h light/8 h dark cycle with 160 μmol m^-2^ s^-1^ of light, at 21 °C and 55 % humidity. Four replicate pots were prepared per treatment. Soil samples (0.5 g each) were removed with a spatula every seven days until day 28, and placed into a Lying Matrix E tube for DNA extraction (FastDNA SPIN Kit for Soil, MP Biomedicals, Solon, USA, see below).

### 2.2 Isolation and identification of cultivable bacteria from soil

Bacterial strains were isolated from the soil after 28 days of treatment with BOA, gramine or quercetin. Soil suspensions (1:1, w/w) were prepared with sterile water, mixed and centrifuged at 2000 x g for 15 min. Aliquots (50 μL) of a 10-fold dilution series were plated on TSB medium (17 g/L casein peptone, 3 g/L soya peptone, 5 g/L NaCl, 2.5 g/L K_2_HPO_4_, 2.5 g/L glucose, 15 g/L agar), YPD medium (10 g/L yeast extract, 20 g/L bacto peptone, 20 g/L glucose, 20 g/L agar), TSM medium (1 g/L K_2_HPO_4_, 0.2 g/L MgSO_4_, 0.1 g/L CaCl_2_, 0.1 g/L NaCl, 0.002 g/L FeCl3, 0.5 g/L KNO3, 0.5 g/L asparagine, 1 g/L mannitol, 15 g/L agar) and malt medium (15 g/L malt extract, 8 g/L yeast extract, 5 g/L glucose, 5 g/L fructose, 10 g/L agar). Plates were incubated for several days at 28°C. Colonies were picked for PCR amplification of the bacterial 16S *rRNA* gene using primers 799F and 1192R which target the V4-V7 region. PCR products were purified by excising the 500 bp bands after electrophoresis in 1.2 % agarose gels. DNA was eluted using the NucleoSpin Gel and PCR Clean-up Kit (Macherey-Nagel). After Sanger sequencing (GATC Biotech, Konstanz, Germany), sequences were searched against the nucleotide database at Genbank using BLASTN. Obtained isolates were stored at −80 °C in microbanks. The 16S rRNA gene sequences of the isolated strains were quality checked and trimmed before they were aligned using the SINA aligner v1.2.11 (Pruesse et al., 2012) and imported into the SSU Ref NR 99 138.1 database. The alignment was manually controlled and a phylogenetic tree was calculated using the maximum-likelihood algorithm PHYML in ARB (Ludwig et al., 2004).

### 2.3 Bacterial community profiling by 16S *rRNA* amplicon sequencing

Total DNA was extracted from the soil using the FastDNA SPIN Kit for Soil (MP Biomedicals, Solon, USA). To the soil in the Lysing Matrix E tube, sodium phosphate buffer and MT buffer were added, and it was homogenized with 2 x 30 s rotations at 6000 rpm in a Precellys tissue homogenizer (Bertin, Frankfurt a. M., Germany). Further extraction steps were performed as described in the manufacturer’s protocol. Genomic DNA from soil was eluted in 60 μL of nuclease free water and purified twice using Agencourt MPure XP beads (Beckman-Coulter, Krefeld, Germany). The purified genomic DNA was used in a two-step PCR procedure to amplify the V4-V7 region of the bacterial 16S *rRNA* gene (primers 799F – 1192R, **Supplementary Table S1**) (Bodenhausen et al., 2013). In the first step, the gene fragment was amplified for each sample in triplicates in a 25 μL reaction volume (Agler et al., 2016). Each reaction contained 0.2 μM of each primer, GoTaq Reaction Buffer (1.5 mM MgCl_2_), PCR Nucleotide Mix (0.2 mM each dNTP), 1.25 U GoTaq G2 DNA polymerase (Promega, Walldorf, Germany), 0.3% bovine serum albumine, 4 ng template DNA and nuclease-free water. The first PCR amplification was performed as follows: 2 min at 94°C; 25 cycles of 15 s at 94°C, 25 s at 55°C, 45 s at 72°C. The products were purified by electrophoresis in 1.2 % agarose gels. The bands at 500 bp were excised and the DNA purified using the NucleoSpin Gel and PCR Clean-up kit (Macherey-Nagel, Düren, Germany). DNA was finally eluted in 20 μL nuclease free water. The purified products were used for a second PCR, which was performed in the same way but with only 15 cycles. In the second PCR, barcoded primers containing Illumina adaptors (B5-F; B5-1 to B5-64; **Supplementary Table S1**) were used. The PCR products were also purified by electrophoresis (see above). The DNA concentration of the purified PCR products was determined by fluorometry (QuantiFluor ONE Dye, Quantus Fluorometer, Promega, Walldorf, Germany), and a sequencing library was created by pooling 30 ng of each amplicon. The library was purified twice with Agencourt MPure XP beads.

### 2.4 Illumina sequencing

The PCR products (100 ng) of the purified sequencing library were used for 2 x 300 bp paired-end Illumina sequencing on a MiSeq sequencer (Illumina, Berlin, Germany). Raw sequence data (lllumina) were analyzed using the Quantitative Insights Into Microbial Ecology software (QIIME 1.9.0) (Caporaso et al., 2010). The taxonomic assignment is based on the Ribosomal Database Project (RDP) using a bootstrap cutoff of 0.5. An amplicon sequence variant (ASVs) table was obtained using the recently revised bacterial taxonomy system (Parks et al., 2018).

### 2.5 Statistics

The ASV table was rarefied to the lowest number of reads (7198 reads) to obtain equal read numbers for all samples. Alpha-diversity indices (Shannon, Chao1, inverse Simpson and Pilou’s evenness) were calculated in R using the package vegan (Oksanen et al., 2019; R Core Team, 2020). Overall differences between groups of samples were assessed using nonparametric Kruskal-Wallis tests while Mann-Whitney tests were applied for individual comparisons, because data were non-normally distributed. A Bonferroni-Holm correction was performed to correct for multiple comparisons. The effects of treatment (control, BOA, gramine and quercetin) and sampling time points (t7, t14, t21 and t28) were evaluated.

A non-metric multidimensional scaling (NMDS) plot based on Bray-Curtis distances was built with Hellinger transformed data. To test for significant differences in the bacterial community composition in dependence on treatment and time, Permutational ANOVA (ADONIS) was performed using the Bray-Curtis distance matrix data, which were derived from Hellinger transformed ASV tables (package ‘vegan’ version 2.5-6 in R).

Responsive taxa significantly enriched in relative abundance due to sample treatment were identified at phylum and genus levels using ANOVA with Tukey-Kramer Post-hoc analyses and a Benjamini-Hochberg false discovery rate correction in STAMP (Parks et al., 2014). A heatmap was generated in R using the packages pheatmap and dplyr (Kolde, 2019; Wickham et al., 2020), which shows the preferential occurrence of the most abundant ASVs (> 0.5% relative abundance) in dependence on treatment and time point. Data were log10 transformed. Dendrograms were calculated using the WPGMA clustering algorithm based on an Euclidean distance matrix derived from relative ASV abundance.

### 2.6 Phospholipid fatty acid analysis

Lipids were extracted from the soil with chloroform/methanol. Phospholipids were isolated by solid phase extraction, fatty acids were converted into their methyl derivatives in the presence of the internal standard tridecanoic acid (13:0) and quantified by gas chromatography-mass spectrometry as described (Siebers et al., 2018).

## 3. Results

### 3.1 Responses of the soil microbiota to plant metabolites

We incubated agricultural soil in open pots with addition of one of the three plant metabolites, BOA, gramine or quercetin (each with a total amount of ~0.5 μmol per g soil) over a period of 28 days. Different alpha diversity indices (Shannon, Chao1, Pilou’s Evenness and inverse Simpson index) as measures for biodiversity, species richness, equal distribution of species within the community and probability for the species identity of two randomly selected individuals, respectively, were calculated (**Figure 1**). All alpha diversity indices showed a significant reduction after BOA and quercetin treatment compared to the control. In contrast, alpha diversity indices of gramine-treated samples were not significantly different from control samples. Changes in all alpha diversity indices over time were inconsistent and rarely significant, indicating no strong alterations during the course of incubation.

**Figure 1.**
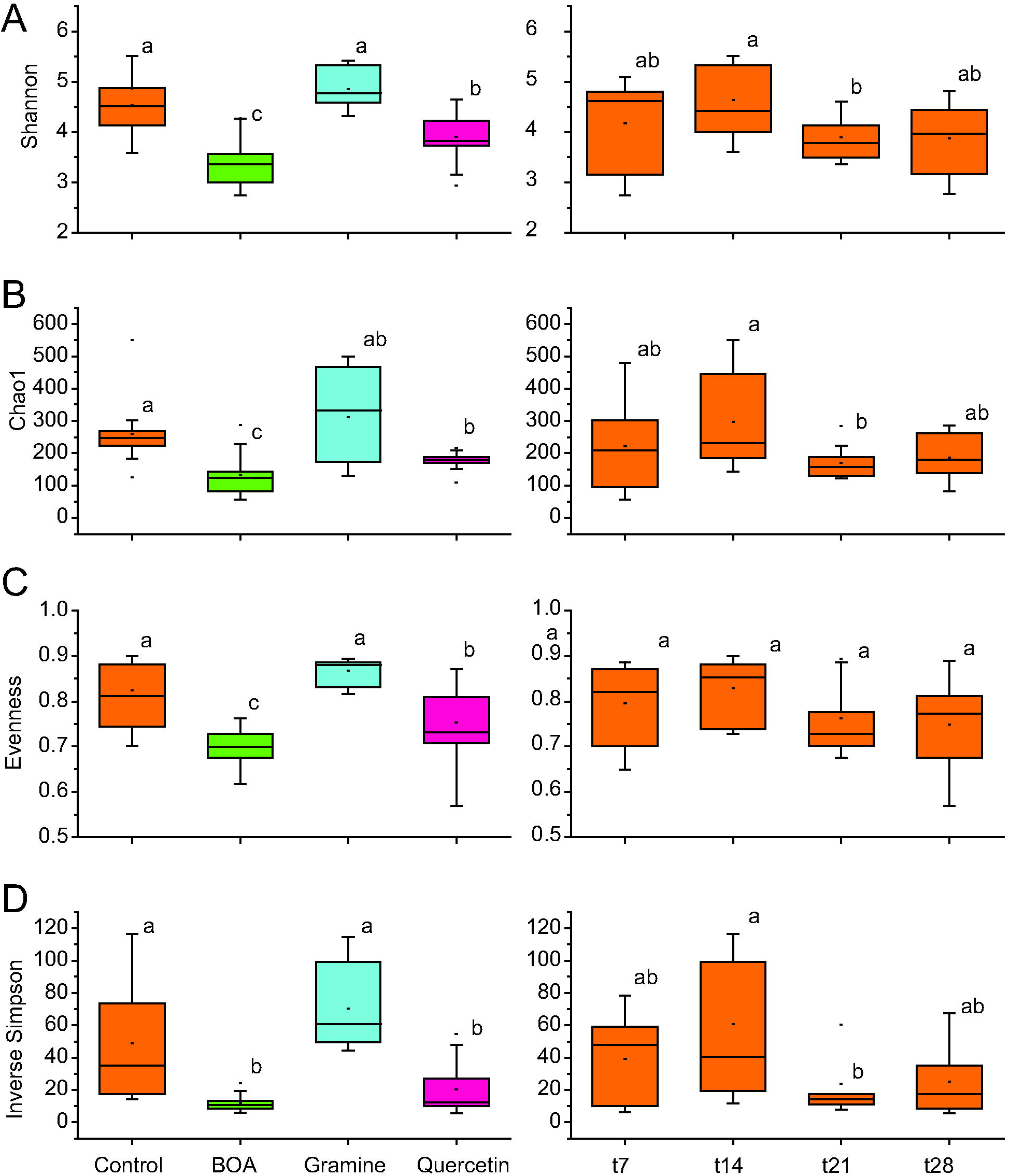
Box plots showing differences in alpha-diversity measures of the soil bacterial communities in dependence on treatments with different plant metabolites (left column) and over time (right column). Boxplots are shown for the Shannon index **(A)**, Chao1 index **(B)**, Pilou’s Evenness index **(C)** and inverse Simpson index **(D)**. Significant differences between groups of samples are indicated by lower-case letters according to Mann-Whitney tests.

Changes in the bacterial community structure were evaluated in NMDS plots (**Figure 2A**). A clear clustering according to treatment was evident, while temporal differences explained less of the variation in community composition between samples. These observations were confirmed by ADONIS with higher R^2^-values for treatment then for time points (treatment, R^2^ = 0.46, *P* = 0.001; time point, R^2^ = 0.10, *P =* 0.001; autocorrelation treatment * time point, R^2^ = 0.18, *P =* 0.001). The samples treated with BOA, gramine or quercetin were located in separate clusters at all time points. For the control samples, the early time points (t7, t14) were separated from the late ones (t21, t28) which partially overlapped with the BOA treated samples. This indicates that the bacterial community structure was strongly altered by gramine and quercetin treatment, while the treatment with BOA resulted in a transient difference in the bacterial community composition compared to the control samples.

**Figure 2.**
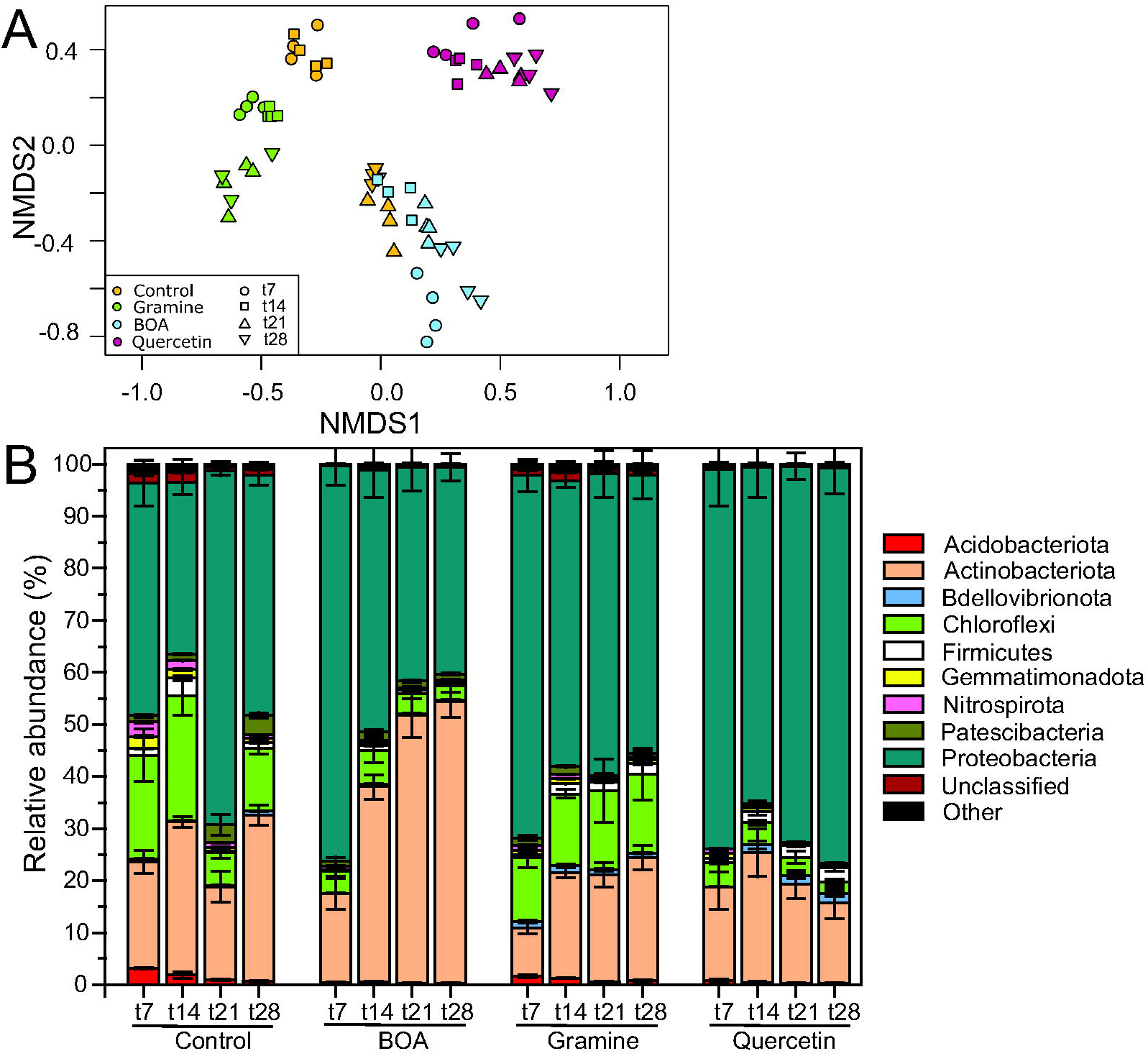
Changes in the soil bacterial community structure after the treatment with BOA, gramine and quercetin and at the different sampling time points. **(A)** Differences in bacterial community structure between samples illustrated in non-metric multidimensional scaling (NMDS) plots. The time points are shown by different symbols, and the color code depicts the different treatments. **(B)** Relative abundance of bacterial phyla in the soil of the control or after treatment with BOA, gramine or quercetin. Low abundant groups with <0.5% of the total reads are summarized as “Other”. (n=4, mean ± SD).

### 3.2 Bacterial taxa are differentially affected by plant metabolites

The phyla of *Actinobacteriota, Chloroflexi* and *Proteobacteria* were most abundant in the control soil samples with mean relative abundances of 31.9%, 12.1% and 46.2% at the time point t28, respectively, while *Acidobacteriota, Bdellovibrionota, Firmicutes, Gemmatimonadota, Nitrospirota, Patescibacteria,* unclassified bacteria and others were much less abundant (all < 2%, **Figure 2B**). Exposure to the plant metabolites revealed clear changes in taxonomic composition. Incubation with BOA resulted in a significant increase in the relative abundances of *Actinobacteriota* up to 54.2% with a concomitant decrease in *Proteobacteria* (39.9%), *Acidobacteriota* (0.2%), *Firmicutes* (0.6%) and *Chloroflexi* (2.7%) at time point t28 compared to the control treatment. Exposure to gramine did not affect relative abundances of *Actinobacteriota* (23.7%) at t28 compared to the control treatment, while *Proteobacteria* and *Bdellivibronota* were significantly more abundant with 53.5% and 0.8%, respectively (**Figure 2B**). Quercetin treatment resulted in significantly lower relative abundances of *Acidobacteriota* (0.2%) and *Chloroflexi* (2.2%), and higher percentages of *Proteobacteria* (76.0%) and *Bdellovibrionota* (1.8%) at t28. All treatment effects were tested with ANOVA.

The treatment with the three plant-derived specialized metabolites also resulted in characteristic changes at the bacterial genus level. Fifty-two genera showed a significant increase in relative abundance in response to the treatment with BOA, gramine or quercetin (all time points aggregated) (**Supplementary Table S2**). The genera with relative abundances > 0.5% at any time point in the control or one of the three treatments were included in a heatmap showing the changes in relative abundances between treatments and time points (**Figure 3**). Clustering of the samples reflected NMDS results with a clear separation of samples according the three treatments with the plant metabolites. Early time points of the control clustered most closely to the gramine treatment, while late control time points were most similar to the BOA treatment.

**Figure 3.**
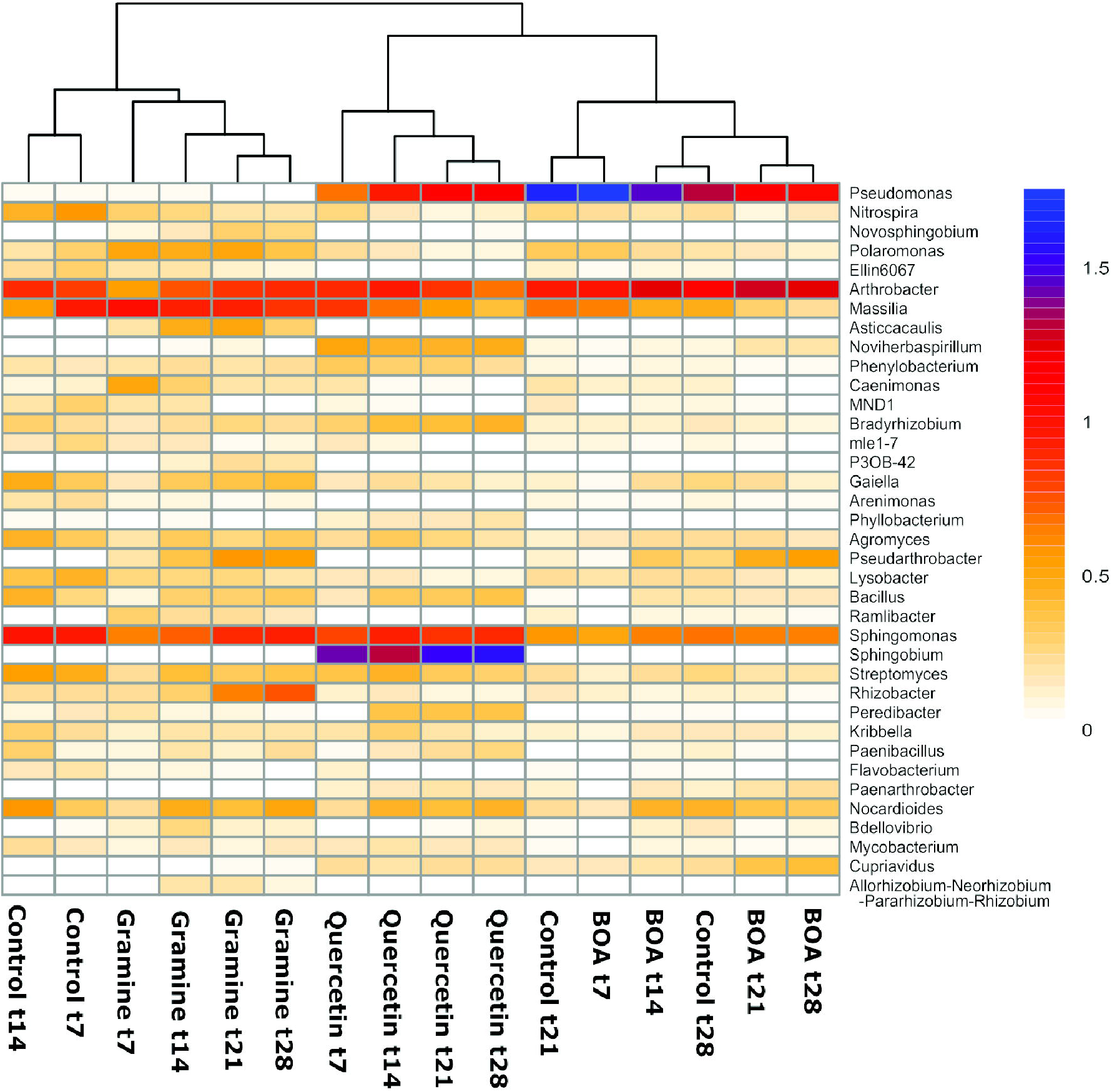
Heatmap showing differences in relative abundances of abundant (>0.5 %) bacterial genera in soil after application of different plant metabolites over time. Differences in relative abundance (log10-transformed) are color-coded with blue colors indicating high relative abundances, and yellow colors indicating low abundances. The dendrogram was derived from WPGMA clustering of an Euclidean distance matrix.

Genera with a significant increase or decrease in relative abundance after treatment with the metabolites are presented in two Venn diagrams (**Figure 4**). Three genera of *Actinobacteriota (Arthrobacter*, *Pseudarthrobacter*, *Paenarthrobacter)* and one of *Proteobacteria (Cupriavidus),* showed an increase in relative abundance after BOA treatment. The relative abundance of *Pseudarthrobacter* was also increased after gramine treatment, while *Cupriavidus* and *Paenarthrobacter* also accumulated after quercetin treatment. In addition, gramine treatment caused the accumulation of 10 different *Proteobacteria* genera, two genera of *Bdellovibronota* and one genus of *Myxococcota.* The genera that increased in relative abundance upon quercetin treatment were mostly *Proteobacteria* (9 genera), *Actinobacteriota* and *Bdellovibrionota*. Many genera decreased in relative abundance, some even below the detection limit, after application of either of the three metabolites.

**Figure 4.**
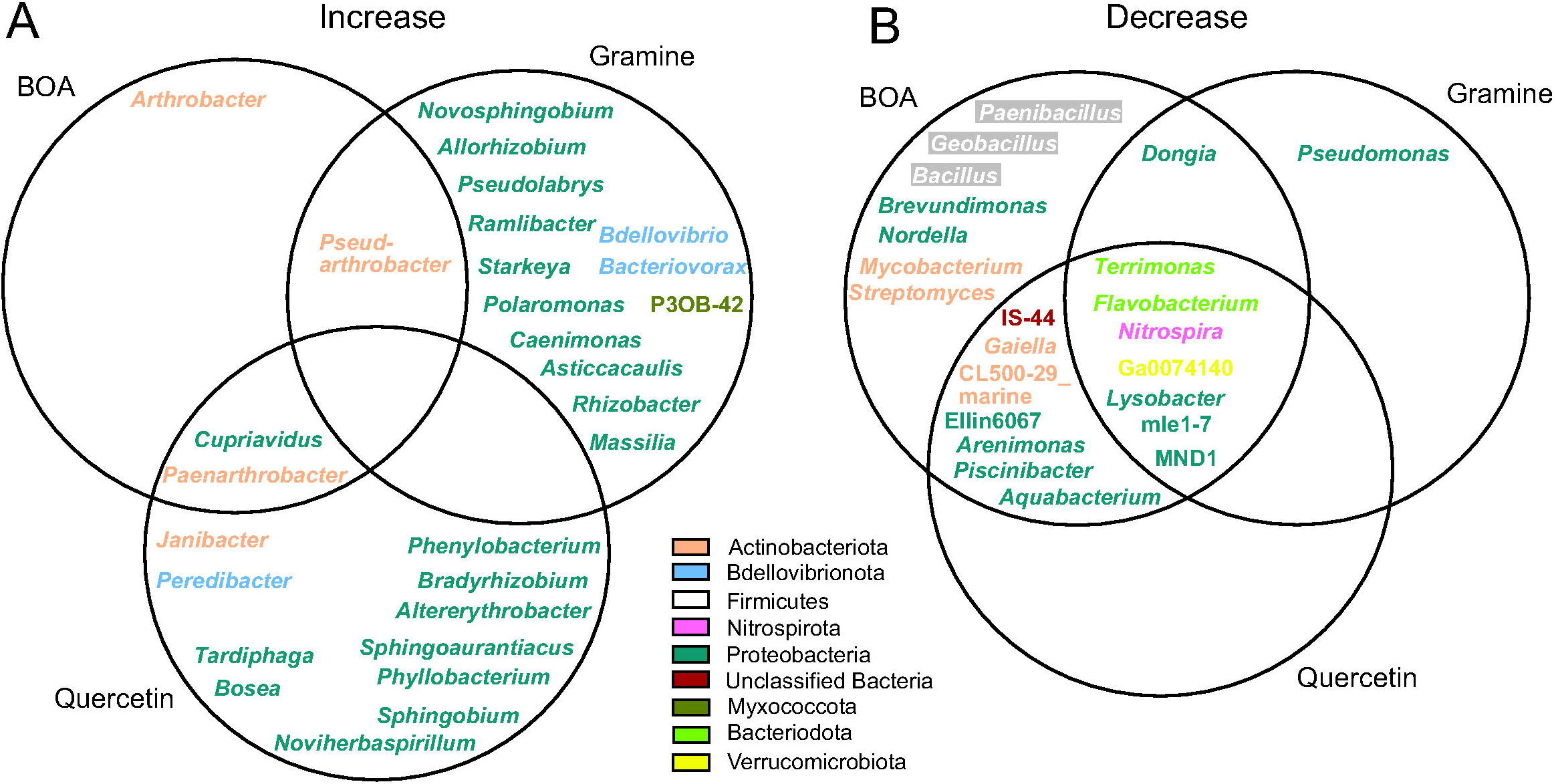
Venn diagrams showing genera with increased or decreased relative abundance after treatment with BOA, gramine or quercetin in comparison to the control. Different colors indicate the assignment to the different phyla.

**Supplementary Figure S1** shows the changes in relative abundances during the metabolite treatment for some selected individual ASVs. A large number of ASVs, e.g. *Massilia,* showed a decrease in relative abundance after the treatment with one or several metabolites, indicating that their growth was limited by the plant metabolites in comparison with other bacteria. Other ASVs like *Arthrobacter* ASV2 or *Pseudarthrobacter,* showed a gradual increase in relative abundance during metabolite treatment, and they were also increased in the control experiment, albeit to a lower extent, suggesting that these bacteria are tolerant to the metabolites. A few ASVs like *Arthrobacter* ASV1 showed a maximum of relative abundance after 14 and 21 days of treatment (**Figure S1A**). Other ASVs accumulated to very high relative abundances at least in one of the treatments, for example *Arthrobacter* ASV3 which was abundant during BOA application, but was also found in the other treatments (**Figure S1B**). A *Pseudomonas* ASV showed high relative abundances of > 20% at day 7 after BOA treatment and then declined, while this ASV also accumulated later in the control samples peaking at day 21. A *Sphingobium* ASV accumulated up to 28 % only during quercetin treatment. Furthermore, an unclassified ASV from the *Micrococcaceae* increased to more than 30% after BOA treatment. The distinct patterns of changes in relative abundances during metabolite treatments of the individual ASVs (**Figure S1**) as compared with the corresponding changes on genus level (**Figures 3, 4**) demonstrate that the ASVs show very diverse responses which are lost in many cases when only condensing the responses of the whole genus.

Taken together, treatment with BOA resulted in the increase in relative abundance of only four genera, three of them belonging to the *Actinobacteriota,* while gramine or quercetin treatment caused an increase in relative abundance of many genera, particularly members of the *Proteobacteria.* On the other hand, BOA treatment caused a decrease in relative abundance of a high number of genera, while some of the relative abundances were also decreased by quercetin or gramine. However, only very few genera were exclusively decreased in abundance by gramine or quercetin alone (**Figure 4**).

### 3.3 Phospholipid fatty acid measurements of soil samples

The analysis of the phospholipid fatty acid (PLFA) content and composition represents a valuable tool to study the microbial diversity in the soil. Phospholipids were extracted and purified from the soil after 28 days of incubation, and PLFA measured by GC-MS after conversion into methyl esters. The total amounts of PLFA were around 0.15 μg/g soil in all samples without significant differences (**Figure 5**). Because PLFA are the major components of cellular membranes, they can serve a measure for the total amount of microbial biomass in the soil. Therefore, the total amount of microbiota in the soil was not affected after the treatment with BOA, gramine or quercetin in comparison with the control. Specific PLFAs are characteristic for bacterial or fungal taxa or for cyanobacteria and eukaryotic algae (Frostegård and Bååth, 1996). The most abundant PLFA detected in the soil samples were the saturated fatty acids 16:0 and 18:0. Furthermore, odd chain (e.g. 15:0, 17:0), methyl-branched (e.g. 15:0iso, 15:0anteiso), monounsaturated (e.g. 16:1Δ9, 16:1Δ7, 18:1Δ11) and cyclopropane (17:0cyclo) fatty acids were identified. Odd chain, methyl-branched and monounsaturated fatty acids are mainly found in bacteria, while cyclopropane fatty acids are produced in bacteria after stress (Kaneda, 1991; Alvarez-Ordóñez et al., 2008). Indeed, we observed increases in the cyclopropane fatty acid 17:0cyclo indicating that the bacteria were subjected to stress after metabolite supplementation. Fatty acids typically found in fungi like oleic acid (18:1Δ9) or diunsaturated fatty acids (e.g.18:1Δ9,12), or in cyanobacteria/algae (triunsaturated fatty acids, 16:3Δ7,10,13; 18:3Δ9,12,15) (Frostegård and Bååth, 1996; Ibekwe and Kennedy, 1998; Kaiser et al., 2010), were of low abundance or absent from the soil. Therefore, the soil was dominated by the presence of bacterial microbiota, with very low amounts of fungi.

**Figure 5.**
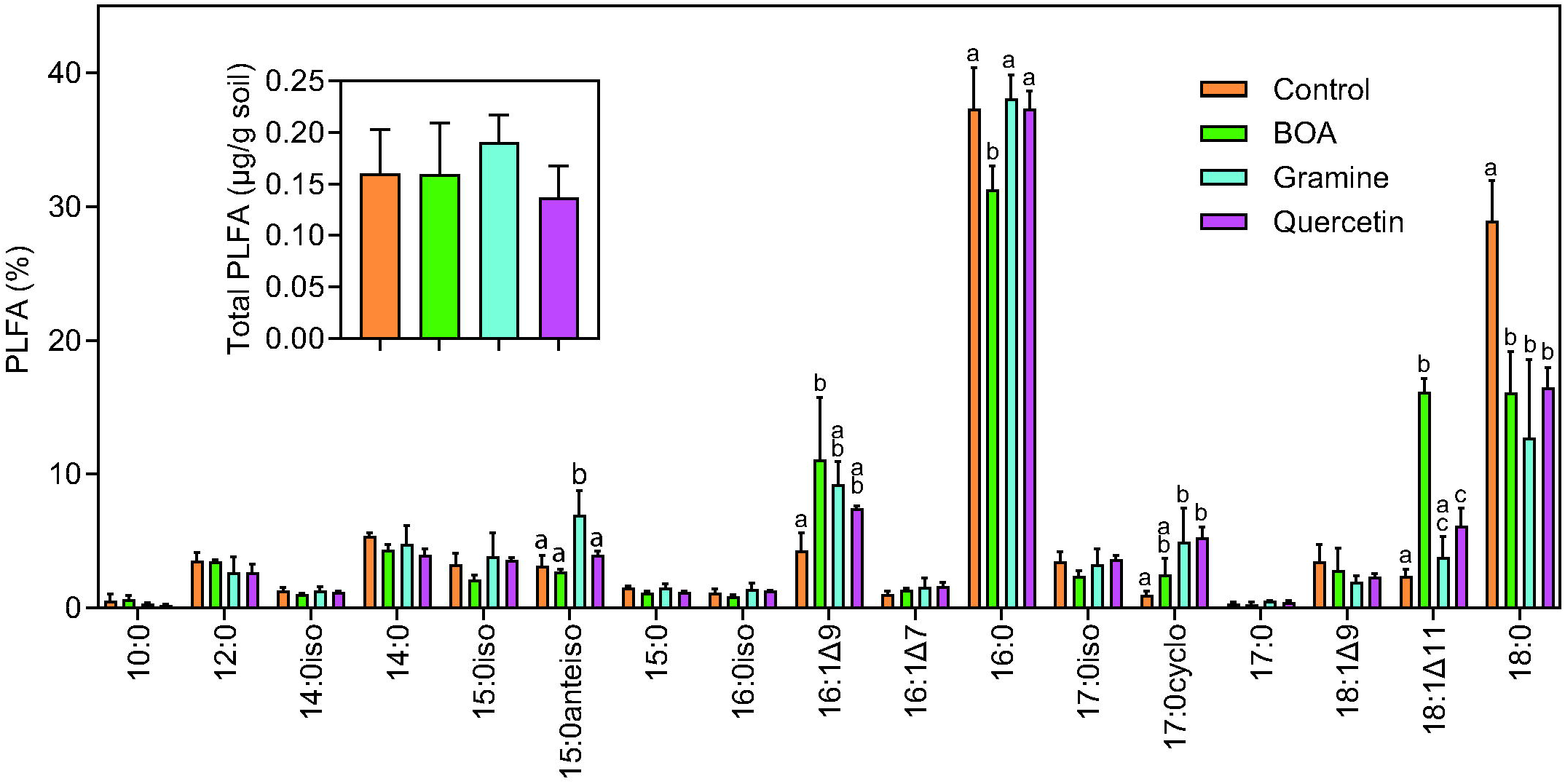
Phospholipid fatty acid (PLFA) analysis of soil samples. Soil samples harvested after 28 days of treatment with BOA, gramine or quercetin were extracted and phospholipid fatty acids determined by GC-MS. Data are shown in %. The inset shows total PLFA in μg per g soil. 10:0, decanoic acid; 12:0, lauric acid; 14:0iso, 11-methyl-tridecanoic acid; 15:0iso, 13-methyl-myristic acid; 15:0anteiso, 12-methyl-myristic acid; 16:0iso, 13-methyl-pentadecanoic acid; 16:1Δ9, palmitoleic acid; 16:1Δ7, Δ7-hexadecenoic acid; 16:0, palmitic acid; 17:0iso, 15-methyl-palmitic acid; 17:0cyclo, 7,8-cyclopropane-palmitic acid; 18:1Δ9, oleic acid; 18:1Δ11, vaccenic acid; 18:0, stearic acid. (ANOVA, post hoc Tukey; p < 0.05; n=3; mean ± SD; different letter indicate significant differences).

### 3.4 Isolation of cultivable strains after treatment of the soil with plant metabolites

To recover microorganisms that can withstand the treatment of the soil with the plant metabolites, microorganisms were isolated from the soil harvested after 28 d of incubation in the presence of BOA, gramine or quercetin and from control soil. The control soil was included to demonstrate that the composition of isolates obtained from metabolite-treated soil differs from that obtained from untreated soil. Isolates were characterized after growing the strains on different media and DNA sequencing of the 16S *rRNA* gene. A total of 107 bacterial strains were isolated from control and metabolite-treated soil (**Supplementary Table S3**). We recovered *Actinobacteriota (Paenarthrobacter*, *Pseudarthrobacter, Arthrobacter, Nocardioides, Mycobacterium, Streptomyces), Proteobacteria (Pseudomonas, Massilia, Cupriavidus, Limnohabitans, Novosphignobium, Sphingobium, Rhizobium, Phyllobacterium)* and *Firmicutes (Bacillus, Paenibacillus)* from control soil and from the treated soil. To assess the relationship between the different strains, the 16S *rRNA* gene sequences were aligned together with the closest type strain sequences, and phylogenetic trees were generated (**Figure 6A, B**). From this analysis it became clear that many cultivable isolates can be divided into groups depending on the treatment. For example, one BOA-dependent *Paenarthrobacter* group, and two distinct groups of the *Pseudarthrobacter* strains from BOA or quercetin treated soil were identified. Further branches encompassed isolates from *Burkholderiales (Massilia, Cupriavidus, Limnohabitans,* isolated from BOA treated soil), a *Novosphingobium* branch dependent on quercetin treatment, *Sphingobium* (BOA dependent) and a *Rhizobiales* group *(Rhizobium, Phyllobacterium,* isolated after BOA treatment) (**Figure 6A, B**).

**Figure 6.**
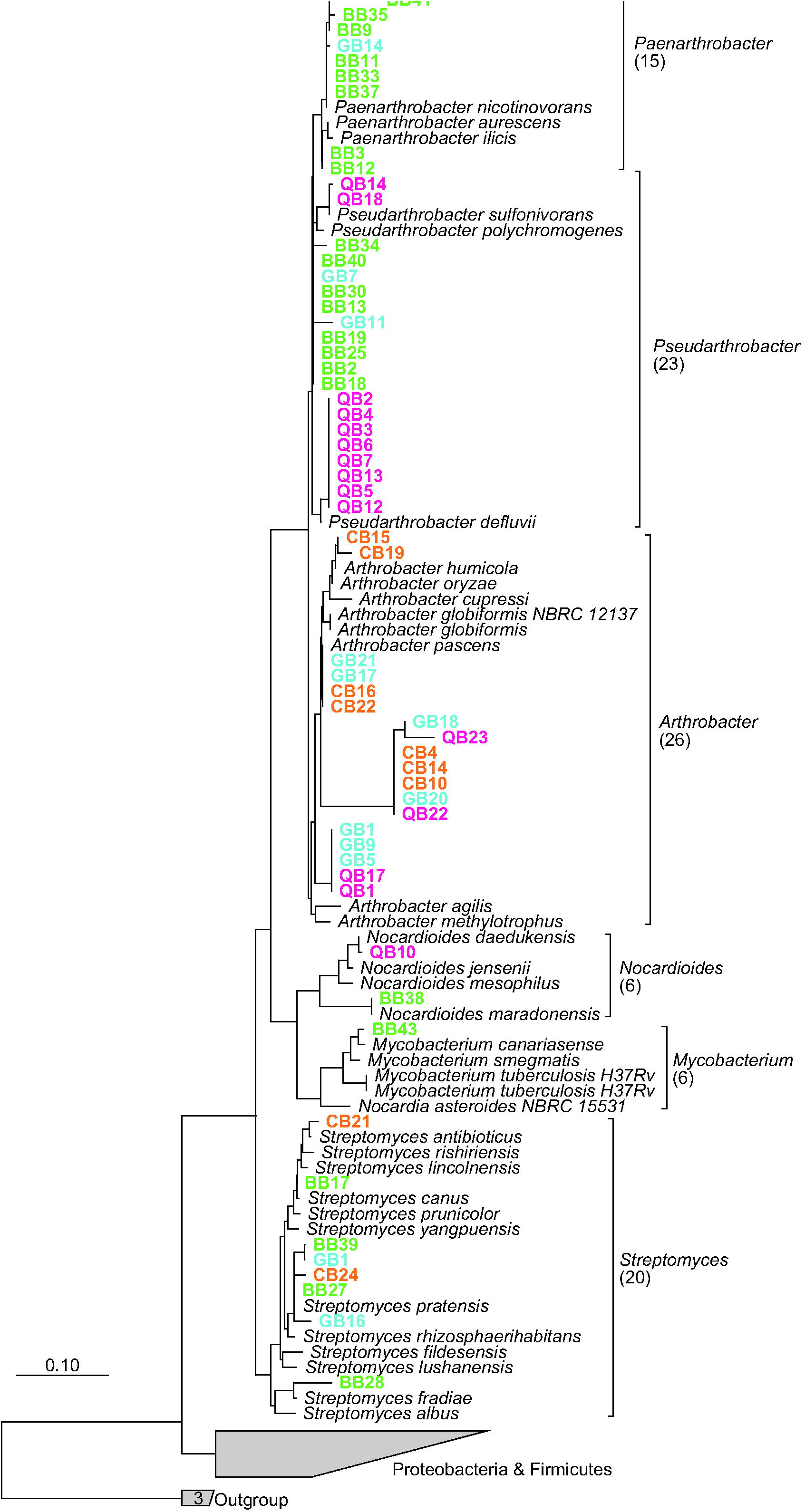
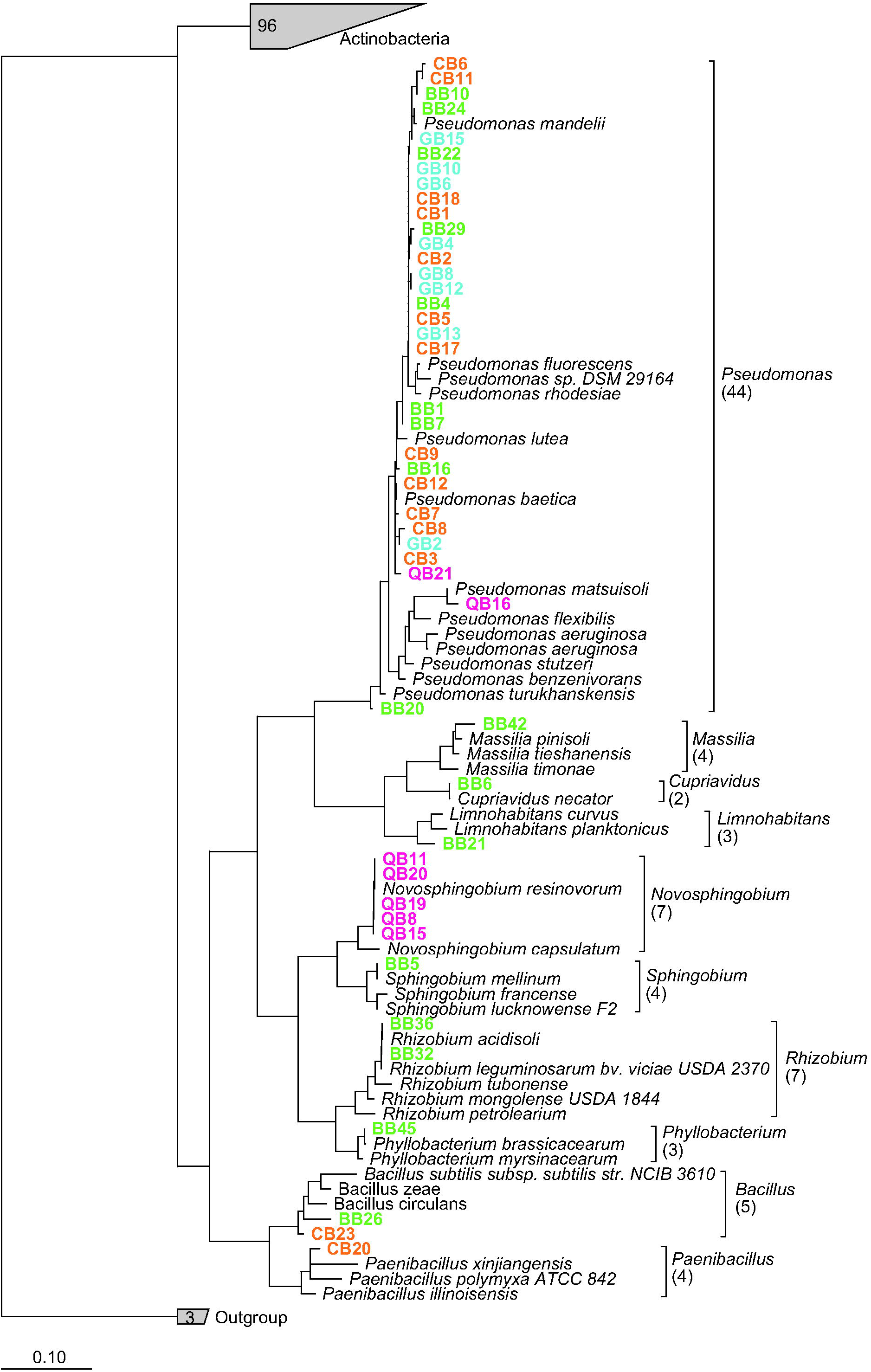
Phylogenetic relationship of bacterial isolates obtained after treatment of the soil with BOA, gramine or quercetin and from control soil. The tree was calculated based on the maximum-likelihood algorithm. Color coding indicates isolates from soils treated with BOA, gramine or quercetin or from control soil. **(A)** *Actinobacteria (Paenarthrobacter*, *Pseudarthrobacter*, *Arthrobacter, Nocardioides, Mycobacterium, Streptomyces).* **(B)** Proteobacteria *(Pseudomonas, Massilia, Cupriavidus, Limnohabitans, Novosphingobium, Sphingobium, Rhizobium)* and Firmicutes *(Bacillus, Paenibacillus)*

## 4. Discussion

### 4.1 General effects of the plant metabolites on soil bacteria

There is accumulating evidence in recent years demonstrating that plant metabolites can change the profile of root-associated microbial communities (Stringlis et al., 2018; Huang et al., 2019; Voges et al., 2019; Harbort et al., 2020). The soil microbiota is crucial for agricultural productivity with respect to e.g. mineral recycling, water regulation and degradation of harmful molecules. A loss of species diversity and reduced abundance in the soil can therefore result in incomplete plant microbiota with reduced functions, resulting in decreased growth, increased susceptibility to diseases, and finally in harvest losses. Recently, Köberl et al. identified edaphic conditions, including soil pH and herbicide usage as important factors for soil microbial community structures in vineyards, orchards and other crop soils (Köberl et al., 2020). Agricultural practices such as continued monocultures without fallow periods can have detrimental effects on the soil microorganisms, and these effects can be caused by the release of plant metabolites into the soil.

In the present study, we focused on the effects of two plant indole metabolites, BOA and gramine, mostly produced by Gramineae species, in comparison with the flavonoid quercetin, on soil bacterial communities. Bacterial 16S *rRNA* gene community profiling revealed that the three metabolites conveyed distinct effects on the community structure of soil-borne bacteria. These alterations are reflected by changes in alpha diversity after BOA and quercetin treatment, while alpha diversity of gramine treatment was not significantly different from control samples (**Figure 1**). The three metabolites caused alterations in taxonomic diversity as revealed in the NMDS plots, while BOA treatment showed in part overlapping data points with the control (**Figure 2A**). These alterations were also observed by changes in relative abundances at the phylum and genus levels (**Figures 2, 3, 4**). Microorganisms able to cope with the plant-derived compounds can remain in a metabolic active stage and may modulate soil parameters, such as pH, to their favor while shaping the species composition of other bacterial groups for optimization of co-existence and cooperation. Adapted microorganisms may group for acting in concert for compound detoxification and degradation. Microorganisms with these abilities obviously remain longer in the soil, even when the crop species change in subsequent cultures. These bacteria might give report about the history of soil usage, and about species particularly adaptable to defined plant metabolites. PLFA analysis demonstrated that the treatment with plant metabolites did not affect the overall content of the microbiota in the soil. Therefore, the alterations in the abundances of ASVs as derived from amplicon sequencing are correlated with absolute changes in the amounts of the corresponding bacteria. Interestingly, the fatty acid composition indicated that the soil was dominated by bacterial microbiota with very little fungal contribution. This result is in accordance with the results of amplicon sequencing because only a low number of fungal sequences were detected.

The effects of BX on bacteria of the rhizosphere were previously studied by growing BX-producing wild type (WT) or deficient (*bx*) maize plants in soil (Hu et al., 2018; Cotton et al., 2019; Kudjordjie et al., 2019). The relative abundance of specific bacterial taxa was decreased in WT compared with the *bx* mutants, and therefore was predominantly negatively affected by the presence of BXs in WT (Cotton et al., 2019; Kudjordjie et al., 2019). While it is known that the microbioms of the rhizosphere and the bulk soil are essentially different because the rhizosphere microbiota represents only a fraction of the bacterial biota in the bulk soil, this finding is in line with results obtained here after direct BOA application to bulk soil, because only four genera were increased, while 22 were decreased (**Figure 4**). Thus, many soil-borne bacteria that do not have rhizosphere competence might not be exposed to physiologically relevant concentrations of BOA by root exudation, but can get in contact with BOA by agricultural practices like crop rotation or mulches. Therefore, the results obtained after BOA application to soil in the present study are relevant for maintaining soil quality. No effects of the plant genotypes of maize WT or *bx* mutants on the alpha diversity of the bacterial microbiota were detected (Hu et al., 2018; Cotton et al., 2019; Kudjordjie et al., 2019), in contrast to the present study where we treated bulk soil directly with BOA causing distinct differences in alpha diversity (**Figure 1**). These contrasting results might be caused by different availability of BXs in the rhizosphere and in the bulk soil. The amounts of metabolites added to the soil in the present work were in the range of the quantities used in previous studies. For example, BXs are released in high amounts (e.g. 0.5 to 5 kg/ha from field-grown rye) into the soil (Barnes and Putnam, 1987; Reberg-Horton et al., 2005). The BX amounts in rye depend on the cultivar and the age of the plants, ranging from 160 to 2000 μg/g dry matter. Understrup and coworkers concluded that amounts of 0.003, 3 and 30 μmol BOA/g soil are naturally reached by root exudation or decaying plant material (Understrup et al., 2005). The biotransformation of 30 μmol BOA/g soil was not completed after 90 days and even later, BOA was still detected. In phytotoxicity experiments, concentrations of 0.1 to 5.0 mM were used to study the effects of BOA on oxidative stress in mung bean (Batish et al., 2006).

Highest amounts of gramine were found in young leaves of wild barley accessions, ranging from 2.032 to 5.290 μmol/g fresh weight, whereas the contents in roots were lower (Maver et al., 2020). Concentrations of 0.5 and 1 mM were employed to measure the phytotoxic effects of gramine on lettuce roots (Maver et al., 2020). Total quercetin in seeds of *Lotus japonicus* amounts to ~1.4 mg/g (~4.6 μmol/g) seeds (Suzuki et al., 2008). Considering that quercetin can be released into the soil, local concentrations of ~0.5 μmol/g soil, might easily be reached. This concentration is in the same range as that of quercetin-glycoside released from white clover (0.5 μmol/g soil), and far below the amounts of luteolin, a flavonoid related to quercetin, released from peanut residues (0.42 μg/g, equivalent to 120 μmol/g soil) (Carlsen et al., 2012; Wang et al., 2018). In the present study, the three metabolites were added to the soil in total amounts ~0.5 μmol/g of soil which is in the range of the naturally occurring concentrations. In fact, HPLC measurements of the contents of BOA, gramine or quercetin in the soil treated for 28 days showed that the metabolites were barely detectable, indicating that their actual concentrations were much lower, presumably because they were bound to soil particles or degraded by microorganisms.

### 4.2 Specific effects on soil genera after application of BOA, gramine or quercetin

Application of BOA, gramine or quercetin to the soil resulted in the enrichment in *Paenarthrobacter*, *Pseudarthrobacter* and *Arthrobacter* (**Figure 4**). These bacteria were also highly enriched in the isolates obtained from treated soil (50 strains of a total of 107 isolates) (**Figure 6A**). The group of *Paenarthrobacter* contains 11 BOA-dependent and one gramine-dependent strain. The first group of *Pseudarthrobacter* isolates encompassed 12 strains, two strains each isolated with quercetin or gramine, and 8 BOA-dependent isolates. The second *Pseudarthrobacter* group contained only strains isolated after quercetin treatment (8 isolates). *Arthrobacter* and related *Actinobacteria* are wide-spread in bulk soil, are resistant to drying and starvation and can therefore live in extreme habitats (Jones and Keddie, 2006). Many *Actinobacteria* strains can degrade pesticides, e.g. a *Paenarthrobacter* isolate was found which is able to metabolize atrazine (Deutch et al., 2018). Therefore, these bacteria harbor detoxification mechanisms and can possibly also metabolize plant metabolites.

Several additional genera related to xenobiotic-degrading strains increased in abundance or were isolated after metabolite treatment, and these genera possibly also harbor pathways to degrade BOA, quercetin or gramin. For example, BOA treatment resulted in increased abundances of two *Nocarioides* and one *Mycobacterium* ASVs. Previously, isolates of *Nocardioides, Variovorax* and *Rhodococcus* were found to degrade xenobiotics like triazine (Martínková et al., 2009; Satsuma, 2010; Satola et al., 2013). Furthermore, specific isolates of *Polaromonas, Ramlibacter* and *Rhizobacter* (increased after gramine treatment) are able to degrade xenobiotics, ginsenoids (triterpene glycosides from ginseng) or natural rubber, respectively (Mattes et al., 2008; Wang et al., 2012; Kasai et al., 2017). In addition, isolates of *Janibacter (Actinobacteriota), Phenylobacterium* and *Sphingobium (Proteobacteria)* (increased with quercetin) can degrade aromatic hydrocarbons, herbicides or pesticides, respectively (Lingens et al., 1985; Cai and Xun, 2002; Zhang et al., 2009), and thus might also be able to metabolize quercetin.

Several genera, including *Cupriavidus, Allorhizobium* and *Bradyrhizobium, Bosea, Tardiaphaga, Phyllobacterium* of the order *Rhizobiales* were increased after treatment with BOA/quercetin, gramine or quercetin, respectively (**Figure 4**). Furthermore, three members of the *Rhizobiales* (two *Rhizobium,* one *Phyllobacterium* strain) were isolated from the soil after BOA treatment (**Figure 6B**). While some members of the Rhizobiales might belong the nonnitrogen fixing Rhizobia, many of which being associated with the root microbiome (Garrido-Oter et al., 2018), others might belong to the nodule-forming Rhizobia which establish symbiotic interactions with legumes, thereby contributing to increase the nitrogen availability in the soil with beneficial effects for the plants that release these metabolites (Vandamme and Coenye, 2004).

Furthermore, strains of *Bdellivibrio, Bacteriovorax,* and Peredibacter showed increased relative abundance after gramine or quercetin treatment, respectively. These bacteria of the Bdellovibrionota are predators feeding on susceptible bacteria (Davidov et al., 2006). The elimination of hazardous or pathogenic bacteria after attraction of *Bdellovibronota* members by gramine or quercetin might have beneficial effects for the plants.

Treatment with gramine or quercetin resulted in the increase in relative abundance of *Novosphingobium* and *Massilia* bacteria, and five *Novosphingobium* isolates were obtained after quercetin treatment. Specific strains of *Novosphingobium* or *Massilia* harbor plant growth promoting properties, e.g. auxin or siderophore production and thus, might also be beneficial for the growth of gramine-or quercetin-producing plants (Ofek et al., 2012; Rangjaroen et al., 2017).

Many *Pseudomonas* isolates were obtained (31 isolates of 107 strains) from control soil or after treatment with BOA, gramine, or quercetin, which might in part be due to the exceptionally high growth rate of *Pseudomonas* strains (**Figure 6B**). Only two *Pseudomonas* strains, QB16 and QB21, were found after quercetin treatment. A large branch contained 29 *Pseudomonas* isolates mostly from control, BOA-treated or gramine-treated soil. The sequence of the ASV BB20 from BOA-treated soil was different from the other *Pseudomonas* sequences and highly similar to that of *P. putida* KT2440, a strain which was attracted to the rhizosphere of 2,4-dihydroxy-1,4-benzoxazin-3-one (DIBOA)-exuding maize roots and harbors plant growth promoting properties because it leads to systemic defense priming (Neal et al., 2012).

BOA application exerted the strongest negative impact on the abundance of bacterial ASVs, with 22 genera being decreased. Many genera were specifically affected by BOA, including three ASVs representing genera of *Firmicutes (Paenibacillus, Geobacillus, Bacillus).* Members of the *Bacillaceae* family were also affected by BX exuded from maize roots (Cotton 2019). The abundance of different *Mycobacterium* and *Streptomyces* was decreased with BOA, in line with previous results which showed that isolates from these genera were sensitive to BXs (Anzai et al., 1960; Atwal et al., 1992; Schütz et al., 2019). The relative abundances of *Flavobacterium, Lysobacter* and *Nitrospira* were decreased by all three metabolites. Previously, quercetin was found to inhibit growth of a *Flavobacterium* strain (Schrader, 2008), and *Flavobacterium and Lysobacter* strains were also affected by BXs exuded from maize roots (Hu et al., 2018; Cotton et al., 2019). *Nitrospira* belongs to the nitrifying bacteria which oxidize ammonia or nitrite to nitrate (Daims and Wagner, 2018). Flavonoids with structures similar to quercetin can inhibit nitrifying bacteria, thereby preventing the oxidation of ammonia in the rhizosphere resulting in a beneficial effect for plant growth (Erickson et al., 2000). The abundance of only one *Pseudomonas* ASV was specifically decreased after gramine treatment. The genus *Pseudomonas* encompasses many soil bacteria, some of which are plant pathogens, like *P. syringae.* It is possible that gramine treatment in the present study decreased the abundance of pathogenic *Pseudomonas* strains, in agreement with previous results which showed that *P. syringae* is sensitive to gramine (Sepulveda and Corcuera, 1990).

## 5. Conclusion

BOA treatment caused the increase in the relative abundance of only four genera, three of them belonging to the *Actinobacteriota,* while 21 ASVs of different phyla were inhibited. Therefore, BOA can be regarded as a broad-spectrum toxin for soil bacteria, with only few genera being able to proliferate. Interestingly, the general impact of the other two metabolites, gramine and quercetin, on soil bacteria, was related, because many more genera were increased after gramine (14) or quercetin (13) treatment, with most genera belonging to the *Proteobacteria.* Almost all bacterial genera inhibited by gramine or quercetin were also inhibited by BOA, suggesting that these strains show a general sensitivity to toxic plant metabolites rather than specifically to gramine or quercetin. Therefore, BOA on the one hand and gramine/quercetin on the other hand reveal different effects on soil bacteria, with BOA showing a broad-spectrum toxic effect preventing the accumulation of many presumably harmful genera, while gramine and quercetin might predominantly exert their function by attracting beneficial genera.

## 6. Contribution to the Field Statement

Plant metabolites can be exuded from the roots or incorporated into the soil from decaying plant material or mulches. Specific metabolites, e.g. glucosinolates produced in rapeseed, can affect the bacterial and fungal community composition in the rhizosphere and the soil. Specific plants produce different metabolites, in some cases in mutually exclusive manner. For example, benzoxazolinone is produced in some barley (Hordeum) species, while others synthesize gramine. The capacity for benzoxazolinone and gramine synthesis is based on the availability of specific genes, but the functional reason for this difference remains unclear. We studied the impact of benzoxazolinone and gramine on soil bacteria, in comparison with a third metabolite, quercetin, which is produced by a large group of plants. Treatment with all three metabolites resulted in strong and specific alterations of soil bacteria composition. The results suggest that benzoxazolinone exerted a broad-spectrum inhibitory effect on many soil bacteria, in contrast to gramine and quercetin which only inhibited a few strains and favored the growth of many, potentially beneficial strains. As a consequence, benzoxazolinone or gramine biosynthesis which have evolved in different barley species, is accompanied with the association with distinct bacterial communities in the soil, presumably after mutual adaptation during evolution.

## 8. Author Contributions

VS, MS and PD conceived the study. KF, PZ, SH, and PS-L contributed to the data analysis of bioinformatics. VS, JC, PZ and SH contributed to the soil sampling. VS and JC performed DNA extractions and PLFA analysis. VS and PD contributed to draft the article. KF, PS-H, CK, MS and PD contributed to critically review and edit the manuscript. All authors contributed to the article and approved the submitted version.

## 10. Acknowledgements

We thank Meike Siebers (University of Düsseldorf) for help with the protocol of PLFA analysis.

## 11. Data Availability Statement

The datasets (amplicon and Sanger sequences) generated for this study can be found in the NCBI Sequence Read Archive (SRA) under the BioProject accession number PRJNA699185.

## 12 Supplementary Material

The Supplementary Material for this article can be found online at http://www.frontiersin.org/.

**Table S1. Oligonucleotides used for PCR amplification and sequencing.**

**Table S2. Bacterial genera identified in the soil after treatment with secondary metabolites.** Sequences of PCR amplicons of the 16S *rRNA* gene were directly obtained from soil DNA. The genera were identified by sequence comparison. The table shows mean relative frequencies and standard deviations in percent, with all time points of one treatment aggregated. Changes to the control are depicted in different colors.

**Table S3. Sequences of 16S *rRNA* amplicons from isolated cultivable bacteria.**

**Figure S1. Alterations in relative abundances of individual ASVs during treatment with plant metabolites.**

The changes in relative abundances of representative ASVs based on amplicon sequencing are shown for the different days of treatments with BOA, gramine or quercetin.

**A**, Representative ASVs showing typical dynamics of responses to the different plant metabolites.

**B**, Four ASVs that were highly abundant at least during one soil treatment.

