## Supplementary material for "Differential impact of plant secondary metabolites on the soil microbiota": Table S1

**Table S1. Oligonucleotides used for PCR amplification and sequencing.**

| Primer | Sequence (5’-3’) | Function |
| --- | --- | --- |
| B5-R1 | ACGACTGCGACTGGCGAACMGGATTAGATACCC | Sequencing primer |
| B5-R2 | CAGCCATTTAGTGTCACGTCATCCCCACCTTCC | Sequencing primer |
| B5-Index | GGAAGGTGGGGATGACGTGACACTAAATGGCTG | Sequencing primer |
| 799F | AACMGGATTAGATACCCKG | PCR1 forward |
| 1192R | ACGTCATCCCCACCTTCC | PCR1 reverse |
| B5-F | AATGATACGGCGACCACCGAGATCTACACGACTGCGACTGGCGAACMGGATTAGATACCCKG | PCR2 forward |
| B5-1 | TCCCTTGTCTCC | PCR2 reverse 1192R Barcode |
| B5-2 | ACGAGACTGATT | PCR2 reverse 1192R Barcode |
| B5-3 | ACCGGTATGTAC | PCR2 reverse 1192R Barcode |
| B5-4 | TGCATACACTGG | PCR2 reverse 1192R Barcode |
| B5-5 | TGGTCAACGATA | PCR2 reverse 1192R Barcode |
| B5-6 | ATCGCACAGTAA | PCR2 reverse 1192R Barcode |
| B5-7 | GTCGTGTAGCCT | PCR2 reverse 1192R Barcode |
| B5-8 | TACAGCGCATAC | PCR2 reverse 1192R Barcode |
| B5-9 | ATCCTTTGGTTC | PCR2 reverse 1192R Barcode |
| B5-10 | AGTCGAACGAGG | PCR2 reverse 1192R Barcode |
| B5-11 | ACCAGTGACTCA | PCR2 reverse 1192R Barcode |
| B5-12 | CCAATACGCCTG | PCR2 reverse 1192R Barcode |
| B5-13 | GCAACACCATCC | PCR2 reverse 1192R Barcode |
| B5-14 | AGTCGTGCACAT | PCR2 reverse 1192R Barcode |
| B5-15 | AGTTACGAGCTA | PCR2 reverse 1192R Barcode |
| B5-16 | TTGCGTTAGCAG | PCR2 reverse 1192R Barcode |
| B5-17 | TACGAGCCCTAA | PCR2 reverse 1192R Barcode |
| B5-18 | TGTCGCAAATAG | PCR2 reverse 1192R Barcode |
| B5-19 | ACAATAGACACC | PCR2 reverse 1192R Barcode |
| B5-20 | TCTCTACCACTC | PCR2 reverse 1192R Barcode |
| B5-21 | CGATCGAACACT | PCR2 reverse 1192R Barcode |
| B5-22 | ATTGCAAGCAAC | PCR2 reverse 1192R Barcode |
| B5-23 | AGCGCTCACATC | PCR2 reverse 1192R Barcode |
| B5-24 | TCGACCAAACAC | PCR2 reverse 1192R Barcode |
| B5-25 | TGTGTTACTCCT | PCR2 reverse 1192R Barcode |
| B5-26 | TGCACAGTCGCT | PCR2 reverse 1192R Barcode |
| B5-27 | TTCTAGAGTGCG | PCR2 reverse 1192R Barcode |
| B5-28 | ACACCTGCGATC | PCR2 reverse 1192R Barcode |
| B5-29 | ATTCCTCTCCAC | PCR2 reverse 1192R Barcode |
| B5-30 | CATCGACGAGTT | PCR2 reverse 1192R Barcode |
| B5-31 | CACCACAGAATC | PCR2 reverse 1192R Barcode |
| B5-32 | GGTCTTAGCACC | PCR2 reverse 1192R Barcode |
| B5-33 | TATCGCGCGATA | PCR2 reverse 1192R Barcode |
| B5-34 | CTCTACGAACAG | PCR2 reverse 1192R Barcode |
| B5-35 | CTCCTCCCTTAC | PCR2 reverse 1192R Barcode |
| B5-36 | CGTGTTATGTGG | PCR2 reverse 1192R Barcode |
| B5-37 | ATTAGCAGCGTA | PCR2 reverse 1192R Barcode |
| B5-38 | CAAGTTTCCGCG | PCR2 reverse 1192R Barcode |
| B5-39 | CCTTGTTCACCT | PCR2 reverse 1192R Barcode |
| B5-40 | AACCAGCAGATT | PCR2 reverse 1192R Barcode |
| B5-41 | CTAGAGCTCCCA | PCR2 reverse 1192R Barcode |
| B5-42 | CACGCAGTCTAC | PCR2 reverse 1192R Barcode |
| B5-43 | ACAAACATGGTC | PCR2 reverse 1192R Barcode |
| B5-44 | TCGAAACATGCA | PCR2 reverse 1192R Barcode |
| B5-45 | TTCCCACCCATT | PCR2 reverse 1192R Barcode |
| B5-46 | AGCAGAACATCT | PCR2 reverse 1192R Barcode |
| B5-47 | GAAACATCCCAC | PCR2 reverse 1192R Barcode |
| B5-48 | CTGTCAGTGACC | PCR2 reverse 1192R Barcode |
| B5-49 | CGGATCTAGTGT | PCR2 reverse 1192R Barcode |
| B5-50 | TTCTCCATCACA | PCR2 reverse 1192R Barcode |
| B5-51 | ATTTAGGACGAC | PCR2 reverse 1192R Barcode |
| B5-52 | GGTTTAACACGC | PCR2 reverse 1192R Barcode |
| B5-53 | AGACAGTAGGAG | PCR2 reverse 1192R Barcode |
| B5-54 | GCAGATTTCCAG | PCR2 reverse 1192R Barcode |
| B5-55 | AGATGATCAGTC | PCR2 reverse 1192R Barcode |
| B5-56 | TATCACCGGCAC | PCR2 reverse 1192R Barcode |
| B5-57 | CCAGATATAGCA | PCR2 reverse 1192R Barcode |
| B5-58 | GGTCTCCTACAG | PCR2 reverse 1192R Barcode |
| B5-59 | ACAGCTCAAACA | PCR2 reverse 1192R Barcode |
| B5-60 | ATAGCGAACTCA | PCR2 reverse 1192R Barcode |
| B5-61 | AACCGCATAAGT | PCR2 reverse 1192R Barcode |
| B5-62 | CTTGAGAAATCG | PCR2 reverse 1192R Barcode |
| B5-63 | CAGTCGTTAAGA | PCR2 reverse 1192R Barcode |
| B5-64 | CTTCCAACTCAT | PCR2 reverse 1192R Barcode |

Note that the PCR2 reverse 1192R barcode primers B5-1 to Bt-64 are composed of the P7 sequence (CAAGCAGAAGACGGCATACGAGAT), the barcode sequence (see Table), the temp/null sequence (CAGCCATTTAGTGTC) and the 1192R sequence (ACGTCATCCCCACCTTCC) (from 5’ to 3’).
