## Supplementary material for "Differential impact of plant secondary metabolites on the soil microbiota": Table S3

**Table S3. Sequences of 16S RNA amplicons from isolated cultivable bacteria.**

**Bacteria isolated from control soil (CB)**

**CB1** *Pseudomonas sp.* UGGGAGCCUUGAGCUCUUAGUGGCGCAGCUAACGCAUUAAGUUGACCGCCUGGGGAGUACGGCCGCAAGGUUAAAACUCAAAUGAAUUGACGGGGGCCCGCACAAGCGGUGGAGCAUGUGGUUUAAUUCGAAGCAACGCGAAGAACCUUACCAGGCCUUGACAUCCAAUGAACUUUCCAGAGAUGGAUUGGUGCCUUCGGGAACAUUGAGACAGGUGCUGCAUGGCUGUCGUCAGCUCGUGUCGUGAGAUGUUGGGUUAAGUCCCGUAACGAGCGCAACCCUUGUCCUUAGUUACCAGCACGUAAUGGUGGGCACUCUAAGGAGACUGCCGGUGA

**CB2** *Pseudomonas sp.*

UGAGCUCUUAGUGGCGCAGCUAACGCAUUAAGUUGACCGCCUGGGGAGUACGGCCGCAAGGUUAAAACUCAAAUGAAUUGACGGGGGCCCGCACAAGCGGUGGAGCAUGUGGUUUAAUUCGAAGCAACGCGAAGAACCUUACCAGGCCUUGACAUCCAAUGAACUUUCCAGAGAUGGAUUGGUGCCUUCGGGAACAUUGAGACAGGUGCUGCAUGGCUGUCGUCAGCUCGUGUCGUGAGAUGUUGGGUUAAGUCCCGUAACGAGCGCAACCCUUGUCCUUAGUUACCAGCACGUAAUGGUGGGCACUCUAAGGAGACUGCCGGUGA

**CB3** *Pseudomonas sp.*

UGAGCUCUUAGUGGCGCAGCUAACGCAUUAAGUUGACCGCCUGGGGAGUACGGCCGCAAGGUUAAAACUCAAAUGAAUUGACGGGGGCCCGCACAAGCGGUGGAGCAUGUGGUUUAAUUCGAAGCAACGCGAAGAACCUUACCAGGCCUUGACAUCCAAUGAACUUUCCAGAGAUGGAUUGGUGCCUUCGGGAACAUUGAGACAGGUGCUGCAUGGCUGUCGUCAGCUCGUGUCGUGAGAUGUUGGGUUAAGUCCCGUAACGAGCGCAACCCUUGUCCUUAGUUACCAGCACGUUAUGGUGGGCACUCUAAGGAGACUGCCGGUGACA

**CB4** *Arthrobacter sp.*

CGUUUUCCGCGCCGUAGCUAACGCAUUAAGUGCCCCGCCUGGGGAGUACGGCCGCAAGGCUAAAACUCAAAGGAAUUGACGGGGGCCCGCACAAGCGGCGGAGCAUGCGGAUUAAUUCGAUGCAACGCGAAGAACCUUACCAAGGCUUGACAUGGACCGGACCGCCGCAGAAAUGUGGUUUCCCCUUUGGGGCCGGUUCACAGGUGGUGCAUGGUUGUCGUCAGCUCGUGUCGUGAGAUGUUGGGUUAAGUCCCGCAACGAGCGCAACCCUCGUUCCAUGUUGCCAGCGCGUAAUGGCGGGGACUCAUGGGAGACUG

**CB5** *Pseudomonas sp.*

AGCCUUGAGCUCUUAGUGGCGCAGCUAACGCAUUAAGUUGACCGCCUGGGGAGUACGGCCGCAAGGUUAAAACUCAAAUGAAUUGACGGGGGCCCGCACAAGCGGUGGAGCAUGUGGUUUAAUUCGAAGCAACGCGAAGAACCUUACCAGGCCUUGACAUCCAAUGAACUUUCCAGAGAUGGAUUGGUGCCUUCGGGAACAUUGAGACAGGUGCUGCAUGGCUGUCGUCAGCUCGUGUCGUGAGAUGUUGGGUUAAGUCCCGUAACGAGCGCAACCCUUGUCCUUAGUUACCAGCACGUAAUGGUGGGCACUCUAAGGAGACUGCCGG

**CB6** *Pseudomonas sp.*

GCCGUUGGGAGCCUUGAGCUCUUAGUGGCGCAGCUAACGCAUUAAGUUGACCGCCUGGGGAGUACGGCCGCAAGGUUAAAACUCAAAUGAAUUGACGGGGGCCCGCACAAGCGGUGGAGCAUGUGGUUUAAUUCGAAGCAACGCGAAGAACCUUACCAGGCCUUGACAUCCAAUGAACUUUCCAGAGAUGGAUGGGUGCCUUCGGGAACAUUGAGACAGGUGCUGCAUGGCUGUCGUCAGCUCGUGUCGUGAGAUGUUGGGUUAAGUCCCGUAACGAGCGCAACCCUUGUCCUUAGUUACCAGCACGUCAUGGUGGGCACUCUAAGGAGACUGCCGGUGA

**CB7** *Pseudomonas sp.*

GCCGUUGGGAGCCUUGAGCUCUUAGUGGCGCAGCUAACGCAUUAAGUUGACCGCCUGGGGAGUACGGCCGCAAGGUUAAAACUCAAAUGAAUUGACGGGGGCCCGCACAAGCGGUGGAGCAUGUGGUUUAAUUCGAAGCAACGCGAAGAACCUUACCAGGCCUUGACAUCCAAUGAACUUUCCAGAGAUGGAUUGGUGCCUUCGGGAACAUUGAGACAGGUGCUGCAUGGCUGUCGUCAGCUCGUGUCGUGAGAUGUUGGGUUAAGUCCCGUAACGAGCGCAACCCUUGUCCUUAGUUACCAGCACGUUAUGGUGGGCACUCUAAGGAGACUGCCGGUGAC

**CB8** *Pseudomonas sp.*

UCUUACUGGCGCAGCcUAACGCAUUAAGUUGACCGCCUGGGGAGUACGGCCGCAAGGUUAAAACUCAAAUGAAUUGACGGGGGCCCGCACAAGCGGUGGAGCAUGUGGUUUAAUUCGAAGCAACGCGAAGAACCUUACCAGGCCUUGACAUCCAAUGAACUUUCCAGAGAUGGAUUGGUGCCUUCGGGAACAUUGAGACAGGUGCUGCAUGGCUGUCGUCAGCUCGUGUCGUGAGAUGUUGGGUUAAGUCCCGUAACGAGCGCAACCCUUGUCCUUAGUUACC

**CB9** *Pseudomonas sp.*

GGAGCCUUGAGCUCUUAGUGGCGCAGCUAACGCAUUAAGUUGACCGCCUGGGGAGUACGGCCGCAAGGUUAAAACUCAAAUGAAUUGACGGGGGCCCGCACAAGCGGUGGAGCAUGUGGUUUAAUUCGAAGCAACGCGAAGAACCUUACCAGGCCUUGACAUCCAAUGAACUUUCCAGAGAUGGAUUGGUGCCUUCGGGAACAUUGAGACAGGUGCUGCAUGGCUGUCGUCAGCUCGUGUCGUGAGAUGUUGGGUUAAGUCCCGUAACGAGCGCAACCCUUGUCCUUAGUUACCAGCACGUUAUGGUGGGCACUCUAAGGAGACUGCCGGUGAC

**CB10** *Arthrobacter sp.*

UCCACGUUUUCCGCGCCGUAGCUAACGCAUUAAGUGCCCCGCCUGGGGAGUACGGCCGCAAGGCUAAAACUCAAAGGAAUUGACGGGGGCCCGCACAAGCGGCGGAGCAUGCGGAUUAAUUCGAUGCAACGCGAAGAACCUUACCAAGGCUUGACAUGGACCGGACCGCCGCAGAAAUGUGGUUUCCCCUUUGGGGCCGGUUCACAGGUGGUGCAUGGUUGUCGUCAGCUCGUGUCGUGAGAUGUUGGGUUAAGUCCCGCAACGAGCGCAACCCUCGUUCCAUGUUGCCAGCGCGUAAUGGCGGGGACUCAUGGGAGACUGCCGG

**CB11** *Pseudomonas sp.*

UAAGCCGUUGGGgAGCCUUGAGCUCUUAGUGGCGCAGCUAACGCAUUAAGUUGACCGCCUGGGGAGUACGGCCGCAAGGUUAAAACUCAAAUGAAUUGACGGGGGCCCGCACAAGCGGUGGAGCAUGUGGUUUAAUUCGAAGCAACGCGAAGAACCUUACCAGGCCUUGACAUCCAAUGAACUUUCCAGAGAUGGAUGGGUGCCUUCGGGAACAUUGAGACAGGUGCUGCAUGGCUGUCGUCAGCUCGUGUCGUGAGAUGUUGGGUUAAGUCCCGUAACGAGCGCAACCCUUGUCCUUAGUUACCAGCACGUCAUGGUGGGCACUCUAAGGAGACUGCCGGUG

**CB12** *Pseudomonas sp.*

GCCGUUGGGAGCCUUGAGCUCUUAGUGGCGCAGCUAACGCAUUAAGUUGACCGCCUGGGGAGUACGGCCGCAAGGUUAAAACUCAAAUGAAUUGACGGGGGCCCGCACAAGCGGUGGAGCAUGUGGUUUAAUUCGAAGCAACGCGAAGAACCUUACCAGGCCUUGACAUCCAAUGAACUUUCCAGAGAUGGAUUGGUGCCUUCGGGAACAUUGAGACAGGUGCUGCAUGGCUGUCGUCAGCUCGUGUCGUGAGAUGUUGGGUUAAGUCCCGUAACGAGCGCAACCCUUGUCCUUAGUUACCAGCACGUUAUGGUGGGCACUCUAAGGAGACUGCCGGUGAC

**CB14** *Arthrobacter sp.*

UCCACGUUUUCCGCGCCGUAGCUAACGCAUUAAGUGCCCCGCCUGGGGAGUACGGCCGCAAGGCUAAAACUCAAAGGAAUUGACGGGGGCCCGCACAAGCGGCGGAGCAUGCGGAUUAAUUCGAUGCAACGCGAAGAACCUUACCAAGGCUUGACAUGGACCGGACCGCCGCAGAAAUGUGGUUUCCCCUUUGGGGCCGGUUCACAGGUGGUGCAUGGUUGUCGUCAGCUCGUGUCGUGAGAUGUUGGGUUAAGUCCCGCAACGAGCGCAACCCUCGUUCCAUGUUGCCAGCGCGUAAUGGCGGGGACUCAUGGGAGACUG

**CB15** *Arthrobacter sp.*

GAAAGCAUGGGGAGCGAACAGGAUUAGAUACCCUGGUAGUCCAUGCCGUAAACGUUGGGCACUAGGUGUGGGGGACAUUCCACGUUUUCCGCGCCGUAGCUAACGCAUUAAGUGCCCCGCCUGGGGAGUACGGCCGCAAGGCUAAAACUCAAAGGAAUUGACGGGGGCCCGCACAAGCGGCGGAGCAUGCGGAUUAAUUCGAUGCAACGCGAAGAACCUUACCAAGGCUUGACAUGGGCCGGACCGGGCUGGAAACAGUCCUUCCCCUUUGGGGCCGGUUCACAGGUGGUGCAUGGUUGUCGUCAGCUCGUGUCGUGAGAUGUUGGGUUAAGUCCCGCAACGAGCGCAACCCUCGUUCCAUGUUGCCAGCGCGUAAUGGC

**CB16** *Arthrobacter sp.*

AAAGCAUGGGGAGCGAACAGGAUUAGAUACCCUGGUAGUCCAUGCCGUAAACGUUGGGCACUAGGUGUGGGGGACAUUCCACGUUUUCCGCGCCGUAGCUAACGCAUUAAGUGCCCCGCCUGGGGAGUACGGCCGCAAGGCUAAAACUCAAAGGAAUUGACGGGGGCCCGCACAAGCGGCGGAGCAUGCGGAUUAAUUCGAUGCAACGCGAAGAACCUUACCAAGGCUUGACAUGGACCGGACCGCCGCAGAAAUGUGGUUUCCCCUUUGGGGCCGGUUCACAGGUGGUGCAUGGUUGUCGUCAGCUCGUGUCGUGAGAUGUUGGGUUAAGUCCCGCAACGAGCGCAACCCUCGUUCCAUGUUGCCAGCGCGUAAUGGCGGG

**CB17** *Pseudomonas sp.*

ACUGAGGUGCGAAAGCGUGGGGAGCAAACAGGAUUAGAUACCCUGGUAGUCCACGCCGUAAACGAUGUCAACUAGCCGUUGGGAGCCUUGAGCUCUUAGUGGCGCAGCUAACGCAUUAAGUUGACCGCCUGGGGAGUACGGCCGCAAGGUUAAAACUCAAAUGAAUUGACGGGGGCCCGCACAAGCGGUGGAGCAUGUGGUUUAAUUCGAAGCAACGCGAAGAACCUUACCAGGCCUUGACAUCCAAUGAACUUUCCAGAGAUGGAUUGGUGCCUUCGGGAACAUUGAGACAGGUGCUGCAUGGCUGUCGUCAGCUCGUGUCGUGAGAUGUUGGGUUAAGUCCCGUAACGAGCGCAACCCUUGUCCUUAGUUACCAGCACGUAAUGGUGG

**CB18** *Pseudomonas sp.*

GCCGUUGGGAGCCUUGAGCUCUUAGUGGCGCAGCUAACGCAUUAAGUUGACCGCCUGGGGAGUACGGCCGCAAGGUUAAAACUCAAAUGAAUUGACGGGGGCCCGCACAAGCGGUGGAGCAUGUGGUUUAAUUCGAAGCAACGCGAAGAACCUUACCAGGCCUUGACAUCCAAUGAACUUUCCAGAGAUGGAUUGGUGCCUUCGGGAACAUUGAGACAGGUGCUGCAUGGCUGUCGUCAGCUCGUGUCGUGAGAUGUUGGGUUAAGUCCCGUAACGAGCGCAACCCUUGUCCUUAGUUACCAGCACGUAAUGGUGGGCACUCUAAGGAGACUGCCGGUGACAAACCGGA

**CB19** *Arthrobacter sp.*

GGUGUGGGGGGACAUUCCACGUUUUCCGCGCCGUAGCUAACGCAUUAAGUGCCCCGCCUGGGGAGUACGGCCGCAAGGCUAAAACUCAAAGGAAUUGACGGGGGCCCGCACAAGCGGCGGAGCAUGCGGAUUAAUUCGAUGCAACGCGAAGAACCUUACCAAGGCUUGACAUGGGCCGGACCGGGCUGGAAACAGUCCUUCCCCUUUGGGGCCGGUUCACAGGUGGUGCAUGGUUGUCGUCAGCUCGUGUCGUGAGAUGUUGGGUUAAGUCCCGCAACGAGCGCAACCCUCGUUCCAUGUUGCCAGCGCGUAAUGGCGGGGACUCAUGGGAGACUGCCGGGGUCAACUCGGA

**CB20** *Paenibacillus sp.*

UUUCAAUACCCUUGGUGCCGAAGUUAACACAUUAAGCAUUCCGCCUGGGGAGUACGGUCGCAAGACUGAAACUCAAAGGAAUUGACGGGGACCCGCACAAGCAGUGCAGUAUGUGGUUUAAUUCGAAGCAACGCGAAGAACCUUACCAGGUCUUGACAUCCCUCUGACCGUCCUAGAGAUAGGGCUUUCCUUCGGGACAGAGGAGACAGGUGGCGCAUGGUUGUCGUCAGCUCGUGUCGUGAGAUGUUGGGUUAAGUCCCGCAACGAGCGCAACCCUUGAUCUUAGUU

**CB21** *Streptomyces sp.*

UCCACGUCGUCGGUGCCGCAGCUAACGCAUUAAGUUCCCCGCCUGGGGAGUACGGCCGCAAGGCUAAAACUCAAAGGAAUUGACGGGGGCCCGCACAAGCAGCGGAGCAUGUGGCUUAAUUCGACGCAACGCGAAGAACCUUACCAAGGCUUGACAUACACCGGAAACGGCCAGAGAUGGUCGCCCCCUUGUGGUCGGUGUACAGGUGGUGCAUGGCUGUCGUCAGCUCGUGUCGUGAGAUGUUGGGUUAAGUCCCGCAACGAGCGCAACCCUUGUUCUGUGUUGCCAGCAUGCCCUUCGGGGUGAUGGGGACUCACAGGAGA

**CB22** *Arthrobacter sp.*

UCCACGUUUUCCGCGCCGUAGCUAACGCAUUAAGUGCCCCGCCUGGGGAGUACGGCCGCAAGGCUAAAACUCAAAGGAAUUGACGGGGGCCCGCACAAGCGGCGGAGCAUGCGGAUUAAUUCGAUGCAACGCGAAGAACCUUACCAAGGCUUGACAUGGACCGGACCGCCGCAGAAAUGUGGUUUCCCCUUUGGGGCCGGUUCACAGGUGGUGCAUGGUUGUCGUCAGCUCGUGUCGUGAGAUGUUGGGUUAAGUCCCGCAACGAGCGCAACCCUCGUUCCAUGUUGCCAGCGCG

**CB23** *Bacillus sp.*

GCUAAGUGUUAGAGGGUUUCCGCCCUUUAGUGCUGCAGCUAACGCAUUAAGCACUCCGCCUGGGGAGUACGGUCGCAAGACUGAAACUCAAAGGAAUUGACGGGGGCCCGCACAAGCGGUGGAGCAUGUGGUUUAAUUCGAAGCAACGCGAAGAACCUUACCAGGUCUUGACAUCCUCUGACAACUCUAGAGAUAGAGCGUUCCCCUUCGGGGGACAGAGUGACAGGUGGUGCAUGGUUGUCGUCAGCUCGUGUCGUGAGAUGUUGGGUUAAGUCCCGCAACGAGCGCAACCCUUGAUCUUAGUUGCCAGCAUUUAGUUGGGCACUCUAAGGUGACUGCCGGUGACAAACCGGA

**CB24** *Streptomyces sp.*

GUUGGCGACAUUCCACGGUCGGUCGGUGCCGCAGCUAACGCAUUAAGUUCCCCGCCUGGGGAGUACGGCCGCAAGGCUAAAACUCAAAGGAAUUGACGGGGGCCCGCACAAGCAGCGGAGCAUGUGGCUUAAUUCGACGCAACGCGAAGAACCUUACCAAGGCUUGACAUAUACCGGAAAGCAUUAGAGAUAGUGCCCCCCUUGUGGUCGGUAUACAGGUGGUGCAUGGCUGUCGUCAGCUCGUGUCGUGAGAUGUUGGGUUAAGUCCCGCAACGAGCGCAACCCUUGUCCUGUGUUGCCAGCAUGCCCUUCGGGGUGAUGGGGACUCACAGGAGA

**Bacteria isolated from BOA treated soil (BB)**

**BB1** *Pseudomonas sp.*

AGCCUUGAGCUCUUAGUGGCGCAGCUAACGCAUUAAGUUGACCGCCUGGGGAGUACGGCCGCAAGGUUAAAACUCAAAUGAAUUGACGGGGGCCCGCACAAGCGGUGGAGCAUGUGGUUUAAUUCGAAGCAACGCGAAGAACCUUACCAGGCCUUGACAUCCAAUGAACUUUCCAGAGAUGGAUUGGUGCCUUCGGGAACAUUGAGACAGGUGCUGCAUGGCUGUCGUCAGCUCGUGUCGUGAGAUGUUGGGUUAAGUCCCGUAACGAGCGCAACCCUUGUCCUUAGUUACCAGCACGUCAUGGUGGGCACUCUAAGGAGACUGCCGGUGACAAACCGGAG

**BB2** *Pseudarthrobacter sp.*

UCCACGUUUUCCGCGCCGUAGCUAACGCAUUAAGUGCCCCGCCUGGGGAGUACGGCCGCAAGGCUAAAACUCAAAGGAAUUGACGGGGGCCCGCACAAGCGGCGGAGCAUGCGGAUUAAUUCGAUGCAACGCGAAGAACCUUACCAAGGCUUGACAUGAACCGGAAAGACCUGGAAACAGGUGCCCCGCUUGCGGUCGGUUUACAGGUGGUGCAUGGUUGUCGUCAGCUCGUGUCGUGAGAUGUUGGGUUAAGUCCCGCAACGAGCGCAACCCUCGUUCUAUGUUGCCAGCACGUGAUGGUGGGGACUCAUAGGAGACUGCCGGGGUCAACUCGGA

**BB3** *Paenarthrobacter sp.*

UCCACGUUUUCCGCGCCGUAGCUAACGCAUUAAGUGCCCCGCCUGGGGAGUACGGCCGCAAGGCUAAAACUCAAAGGAAUUGACGGGGGCCCGCACAAGCGGCGGAGCAUGCGGAUUAAUUCGAUGCAACGCGAAGAACCUUACCAAGGCUUGACAUGAACCGGAAAGACCUGGAAACAGGUGCCCCGCUUGCGGUCGGUUUACAGGUGGUGCAUGGUUGUCGUCAGCUCGUGUCGUGAGAUGUUGGGUUAAGUCCCGCAACGAGCGCAACCCUCGUUCUAUGUUGCCAGCGCGUUAUGGCGGGGACUCAUAGGAGACUGCCGGGGUCAACUCGG

**BB4** *Pseudomonas sp.*

GGAGCCUUGAGCUCUUAGUGGCGCAGCUAACGCAUUAAGUUGACCGCCUGGGGAGUACGGCCGCAAGGUUAAAACUCAAAUGAAUUGACGGGGGCCCGCACAAGCGGUGGAGCAUGUGGUUUAAUUCGAAGCAACGCGAAGAACCUUACCAGGCCUUGACAUCCAAUGAACUUUCCAGAGAUGGAUUGGUGCCUUCGGGAACAUUGAGACAGGUGCUGCAUGGCUGUCGUCAGCUCGUGUCGUGAGAUGUUGGGUUAAGUCCCGUAACGAGCGCAACCCUUGUCCUUAGUUACCAGCACGUAAUGGUGGGCACUCUAAGGAGACUGCCGGUGACAAACCGGA

**BB5** *Sphingobium sp.*

UGCUGGGGUGCAUGGCAUUUCAGUGGCGCAGCUAACGCAUUAAGUUAUCCGCCUGGGGAGUACGGUCGCAAGAUUAAAACUCAAAGGAAUUGACGGGGGCCUGCACAAGCGGUGGAGCAUGUGGUUUAAUUCGAAGCAACGCGCAGAACCUUACCAACGUUUGACAUCCCUAUCGCGGAUCGUGGAGACACUUUCCUUCAGUUCGGCUGGAUAGGUGACAGGUGCUGCAUGGCUGUCGUCAGCUCGUGUCGUGAGAUGUUGGGUUAAGUCCCGCAACGAGCGCAACCCUCGACUUUAGUUGCCAUCAUUUAGUUGGGUACUCUAAAGUAACCGCCGGUGAUAAGCCGGA

**BB6** *Cupriavidus sp.*

UCUUCAGUAACGUAGCUAACGCGUGAAGUUGACCGCCUGGGGAGUACGGUCGCAAGAUUAAAACUCAAAGGAAUUGACGGGGACCCGCACAAGCGGUGGAUGAUGUGGAUUAAUUCGAUGCAACGCGAAAAACCUUACCUACCCUUGACAUGCCACUAACGAAGCAGAGAUGCAUUAGGUGCCCGAAAGGGAAAGUGGACACAGGUGCUGCAUGGCUGUCGUCAGCUCGUGUCGUGAGAUGUUGGGUUAAGUCCCGCAACGAGCGCAACCCUUGUCUCUAGUUGCUACGAAAGGGCACUCUAGAGAGACUGCCGGUGACAAACCGGA

**BB7** *Pseudomonas sp.*

AGCCUUGAGCUCUUAGUGGCGCAGCUAACGCAUUAAGUUGACCGCCUGGGGAGUACGGCCGCAAGGUUAAAACUCAAAUGAAUUGACGGGGGCCCGCACAAGCGGUGGAGCAUGUGGUUUAAUUCGAAGCAACGCGAAGAACCUUACCAGGCCUUGACAUCCAAUGAACUUUCCAGAGAUGGAUUGGUGCCUUCGGGAACAUUGAGACAGGUGCUGCAUGGCUGUCGUCAGCUCGUGUCGUGAGAUGUUGGGUUAAGUCCCGUAACGAGCGCAACCCUUGUCCUUAGUUACCAGCACGUCAUGGUGGGCACUCUAAGGAGACUGCCGGUGACAAACCGGA

**BB9** *Paenarthrobacter sp.*

ACAUUCCACGUUUUCCGCGCCGUAGCUAACGCAUUAAGUGCCCCGCCUGGGGAGUACGGCCGCAAGGCUAAAACUCAAAGGAAUUGACGGGGGCCCGCACAAGCGGCGGAGCAUGCGGAUUAAUUCGAUGCAACGCGAAGAACCUUACCAAGGCUUGACAUGAACCGGAAAGACCUGGAAACAGGUGCCCCGCUUGCGGUCGGUUUACAGGUGGUGCAUGGUUGUCGUCAGCUCGUGUCGUGAGAUGUUGGGUUAAGUCCCGCAACGAGCGCAACCCUCGUUCUAUGUUGCCAGCGGUUCGGCCGGGGACUCAUAGGAGACUGCCGGGGUC

**BB10** *Pseudomonas sp.*

CCGUUGGGAGCCUUGAGCUCUUAGUGGCGCAGCUAACGCAUUAAGUUGACCGCCUGGGGAGUACGGCCGCAAGGUUAAAACUCAAAUGAAUUGACGGGGGCCCGCACAAGCGGUGGAGCAUGUGGUUUAAUUCGAAGCAACGCGAAGAACCUUACCAGGCCUUGACAUCCAAUGAACUUUCCAGAGAUGGAUUGGUGCCUUCGGGAACAUUGAGACAGGUGCUGCAUGGCUGUCGUCAGCUCGUGUCGUGAGAUGUUGGGUUAAGUCCCGUAACGAGCGCAACCCUUGUCCUUAGUUACCAGCACGUCAUGGUGGGCACUCUAAGGAGACUGCCGGUGACAAACCGGA

**BB11** *Paenarthrobacter sp.*

GUGUGGGGGACAUUCCACGUUUUCCGCGCCGUAGCUAACGCAUUAAGUGCCCCGCCUGGGGAGUACGGCCGCAAGGCUAAAACUCAAAGGAAUUGACGGGGGCCCGCACAAGCGGCGGAGCAUGCGGAUUAAUUCGAUGCAACGCGAAGAACCUUACCAAGGCUUGACAUGAACCGGAAAGACCUGGAAACAGGUGCCCCGCUUGCGGUCGGUUUACAGGUGGUGCAUGGUUGUCGUCAGCUCGUGUCGUGAGAUGUUGGGUUAAGUCCCGCAACGAGCGCAACCCUCGUUCUAUGUUGCCAGCGGUUCGGCCGGGGACUCAUAGGAGACUGCCGGGGUCAACUCGGA

**BB12** *Paenarthrobacter sp.*

GUGGGGGACAUUCCACGUUUUCCGCGCCGUAGCUAACGCAUUAAGUGCCCCGCCUGGGGAGUACGGCCGCAAGGCUAAAACUCAAAGGAAUUGACGGGGGCCCGCACAAGCGGCGGAGCAUGCGGAUUAAUUCGAUGCAACGCGAAGAACCUUACCAAGGCUUGACAUGAACCGGAAAGACCUGGAAACAGGUGCCCCGCUUGCGGUCGGUUUACAGGUGGUGCAUGGUUGUCGUCAGCUCGUGUCGUGAGAUGUUGGGUUAAGUCCCGCAACGAGCGCAACCCUCGUUCUAUGUUGCCAGCGCGUUAUGGCGGGGACUCAUAGGAGACUGCCGGGGUCAACUC

**BB13** *Pseudarthrobacter sp.*

UUCCACGUUUUCCGCGCCGUAGCUAACGCAUUAAGUGCCCCGCCUGGGGAGUACGGCCGCAAGGCUAAAACUCAAAGGAAUUGACGGGGGCCCGCACAAGCGGCGGAGCAUGCGGAUUAAUUCGAUGCAACGCGAAGAACCUUACCAAGGCUUGACAUGAACCGGAAAGACCUGGAAACAGGUGCCCCGCUUGCGGUCGGUUUACAGGUGGUGCAUGGUUGUCGUCAGCUCGUGUCGUGAGAUGUUGGGUUAAGUCCCGCAACGAGCGCAACCCUCGUUCUAUGUUGCCAGCACGUGAUGGUGGGGACUCAUAGGAGACUGCCGGGGUCAACUC

**BB15** *Paenarthrobacter sp.*

GUGUGGGGGGACAUUCCACGUUUUCCGCGCCGUAGCUAACGCAUUAAGUGCCCCGCCUGGGGAGUACGGCCGCAAGGCUAAAACUCAAAGGAAUUGACGGGGGCCCGCACAAGCGGCGGAGCAUGCGGAUUAAUUCGAUGCAACGCGAAGAACCUUACCAAGGCUUGACAUGAACCGGAAAGACCUGGAAACAGGUGCCCCGCUUGCGGUCGGUUUACAGGUGGUGCAUGGUUGUCGUCAGCUCGUGUCGUGAGAUGUUGGGUUAAGUCCCGCAACGAGCGCAACCCUCGUUCUAUGUUGCCAGCGGUUCGGCCGGGGACUCAUAGGAGACUGCCGGGGUCAACUCGGA

**BB16** *Pseudomonas sp.*

GUUGGGAGCCUUGAGCUCUUAGUGGCGCAGCUAACGCAUUAAGUUGACCGCCUGGGGAGUACGGCCGCAAGGUUAAAACUCAAAUGAAUUGACGGGGGCCCGCACAAGCGGUGGAGCAUGUGGUUUAAUUCGAAGCAACGCGAAGAACCUUACCAGGCCUUGACAUCCAAUGAACUUUCCAGAGAUGGAUUGGUGCCUUCGGGAGCAUUGAGACAGGUGCUGCAUGGCUGUCGUCAGCUCGUGUCGUGAGAUGUUGGGUUAAGUCCCGUAACGAGCGCAACCCUUGUCCUUAGUUACCAGCACGUUAUGGUGGGCACUCUAAGGAGACUGCCGGUGACAAACCGGA

**BB17** *Streptomyces sp.*

CGACAUUCCACGUCGUCGGUGCCGCAGCUAACGCAUUAAGUUCCCCGCCUGGGGAGUACGGCCGCAAGGCUAAAACUCAAAGGAAUUGACGGGGGCCCGCACAAGCAGCGGAGCAUGUGGCUUAAUUCGACGCAACGCGAAGAACCUUACCAAGGCUUGACAUACACCGGAAAGCAUCAGAGAUGGUGCCCCCCUUGUGGUCGGUGUACAGGUGGUGCAUGGCUGUCGUCAGCUCGUGUCGUGAGAUGUUGGGUUAAGUCCCGCAACGAGCGCAACCCUUGUUCUGUGUUGCCAGCAUGCCCUUCGGGGUGAUGGGGACUCACAGGAGACCGCCGGGGUCAACUCG

**BB18** *Pseudarthrobacter sp.*

GGACAUUCCACGUUUUCCGCGCCGUAGCUAACGCAUUAAGUGCCCCGCCUGGGGAGUACGGCCGCAAGGCUAAAACUCAAAGGAAUUGACGGGGGCCCGCACAAGCGGCGGAGCAUGCGGAUUAAUUCGAUGCAACGCGAAGAACCUUACCAAGGCUUGACAUGAACCGGAAAGACCUGGAAACAGGUGCCCCGCUUGCGGUCGGUUUACAGGUGGUGCAUGGUUGUCGUCAGCUCGUGUCGUGAGAUGUUGGGUUAAGUCCCGCAACGAGCGCAACCCUCGUUCUAUGUUGCCAGCACGUGAUGGUGGGGACUCAUAGGAGACUGCCGGGGUCAACUCGGA

**BB19** *Pseudarthrobacter sp.*

UUCCACGUUUUCCGCGCCGUAGCUAACGCAUUAAGUGCCCCGCCUGGGGAGUACGGCCGCAAGGCUAAAACUCAAAGGAAUUGACGGGGGCCCGCACAAGCGGCGGAGCAUGCGGAUUAAUUCGAUGCAACGCGAAGAACCUUACCAAGGCUUGACAUGAACCGGAAAGACCUGGAAACAGGUGCCCCGCUUGCGGUCGGUUUACAGGUGGUGCAUGGUUGUCGUCAGCUCGUGUCGUGAGAUGUUGGGUUAAGUCCCGCAACGAGCGCAACCCUCGUUCUAUGUUGCCAGCACGUGAUGGUGGGGACUCAUAGGAGACUGCCGGGGUCAACUCGGA

**BB20** *Pseudomonas sp.*

AGUGGCGCAGCUAACGCAUUAAGUUAGUACCGCCUGGGGAGUACGGCCGCAAGAUUAAAACUCAAAUGAAUUGACGGGGGCCCGCACAAGCGGUGGAGCAUGUGGUUUAAUUCGAAGCAACGCGAAGAACCUUACCUGGCCUUGACAUGCUGAGAACUUUCCAGAGAUGGAUUGGUGCCUUCGGGAACUCAAACACAGGUGCUGCAUGGCUGUCGUCAGCUCGUGUCGUGAGAUGUUGGGUUAAGUCCCGUAACGAGCGCAACCCUUGUCCUUAGUUACCAGCACGUUAUGGUGGGCACUCUAAGGAGACUGCCGGUGACAAACCGGA

**BB21** *Limnohabitans sp.*

GGGUCUUCCUGACUCAGUAACGAAGCUAACGCGUGAAGUUGACCGCCUGGNUNUUACGGCCGCAAGGUUGAAACUCAAAGGAAUUGACGGGGACCCGCACAAGCGGUGGAUGAUGUGGUUUAAUUCGAUGCAACGCGAAAAACCUUACCCACCUUUGACAUGUACGGAAUUCGCCAGAGAUGGCUUAGUGCUCGAAAGAGAACCGUAACACAGGUGCUGCAUGGCUGUCGUCAGCUCGUGUCGUGAGAUGUUGGGUUAAGUCCCGCAACGAGCGCAACCCUUGUCAUUAGUUGCUACAUUUAGUUGGGCACUCUAAUGAGACUGCCGGUGACAAACCGGA

**BB22** *Pseudomonas sp.*

CCGUUGGGAGCCUUGAGCUCUUAGUGGCGCAGCUAACGCAUUAAGUUGACCGCCUGGGGAGUACGGCCGCAAGGUUAAAACUCAAAUGAAUUGACGGGGGCCCGCACAAGCGGUGGAGCAUGUGGUUUAAUUCGAAGCAACGCGAAGAACCUUACCAGGCCUUGACAUCCAAUGAACUUUCCAGAGAUGGAUUGGUGCCUUCGGGAACAUUGAGACAGGUGCUGCAUGGCUGUCGUCAGCUCGUGUCGUGAGAUGUUGGGUUAAGUCCCGUAACGAGCGCAACCCUUGUCCUUAGUUACCAGCACGUAAUGGUGGGCACUCUAAGGAGACUGCCGGUGACAAACCGGA

**BB23** *Paenarthrobacter sp.*

UGUGGGGGGACAUUCCACGUUUUCCGCGCCGUAGCUAACGCAUUAAGUGCCCCGCCUGGGGAGUACGGCCGCAAGGCUAAAACUCAAAGGAAUUGACGGGGGCCCGCACAAGCGGCGGAGCAUGCGGAUUAAUUCGAUGCAACGCGAAGAACCUUACCAAGGCUUGACAUGAACCGGAAAGACCUGGAAACAGGUGCCCCGCUUGCGGUCGGUUUACAGGUGGUGCAUGGUUGUCGUCAGCUCGUGUCGUGAGAUGUUGGGUUAAGUCCCGCAACGAGCGCAACCCUCGUUCUAUGUUGCCAGCGGUUCGGCCGGGGACUCAUAGGAGACUGCCGGGGUCAACUCGGA

**BB24** *Pseudomonas sp.*

CCGUUGGGGAGCCUUGAGCUCUUAGUGGCGCAGCUAACGCAUUAAGUUGACCGCCUGGGGAGUACGGCCGCAAGGUUAAAACUCAAAUGAAUUGACGGGGGCCCGCACAAGCGGUGGAGCAUGUGGUUUAAUUCGAAGCAACGCGAAGAACCUUACCAGGCCUUGACAUCCAAUGAACUUUCCAGAGAUGGAUUGGUGCCUUCGGGAACAUUGAGACAGGUGCUGCAUGGCUGUCGUCAGCUCGUGUCGUGAGAUGUUGGGUUAAGUCCCGUAACGAGCGCAACCCUUGUCCUUAGUUACCAGCACGUAAUGGUGGGCACUCUAAGGAGACUGCCGGUGACAAACCGGA

**BB25** *Pseudarthrobacter sp.*

UCCACGUUUUCCGCGCCGUAGCUAACGCAUUAAGUGCCCCGCCUGGGGAGUACGGCCGCAAGGCUAAAACUCAAAGGAAUUGACGGGGGCCCGCACAAGCGGCGGAGCAUGCGGAUUAAUUCGAUGCAACGCGAAGAACCUUACCAAGGCUUGACAUGAACCGGAAAGACCUGGAAACAGGUGCCCCGCUUGCGGUCGGUUUACAGGUGGUGCAUGGUUGUCGUCAGCUCGUGUCGUGAGAUGUUGGGUUAAGUCCCGCAACGAGCGCAACCCUCGUUCUAUGUUGCCAGCACGUGAUGGUGGGGACUCAUAGGAGACUGCCGGGGUCAACUCGGA

**BB26** *Bacillus sp.*

UGUUAGAGGGUUUCCGCCCUUUAGUGCUGAAGUUAACGCAUUAAGCACUCCGCCUGGGGAGUACGGCCGCAAGGCUGAAACUCAAAGGAAUUGACGGGGGCCCGCACAAGCGGUGGAGCAUGUGGUUUAAUUCGAAGCAACGCGAAGAACCUUACCAGGUCUUGACAUCCUCUGAAAACCCUAGAGAUAGGGCUUCUCCUUCGGGAGCAGAGUGACAGGUGGUGCAUGGUUGUCGUCAGCUCGUGUCGUGAGAUGUUGGGUUAAGUCCCGCAACGAGCGCAACCCUUGAUCUUAGUUGCCAUCAUUAAGUUGGGCACUCUAAGGUGACUGCCGGUGACAAACCGGA

**BB27** *Streptomyces sp.*

GGUGUUGGCGACAUUCCACGUCGUCGGUGCCGCAGCUAACGCAUUAAGUUCCCCGCCUGGGGAGUACGGCCGCAAGGCUAAAACUCAAAGGAAUUGACGGGGGCCCGCACAAGCAGCGGAGCAUGUGGCUUAAUUCGACGCAACGCGAAGAACCUUACCAAGGCUUGACAUAUACCGGAAAGCAUCAGAGAUGGUGCCCCCCUUGUGGUCGGUAUACAGGUGGUGCAUGGCUGUCGUCAGCUCGUGUCGUGAGAUGUUGGGUUAAGUCCCGCAACGAGCGCAACCCUUGUUCUGUGUUGCCAGCAUGCCCUUCGGGGUGAUGGGGACUCACAGGAGACUGCCGGGGUCAACUCGGA

**BB28** *Streptomyces sp.*

CCCGUCUGGGAGAGUACGGCCGCAAGGCUAACACUCAAAGGAAUUGACGGGGGCCCGCACAAGCGGCGGAGCGUGUGGCUUAAUUCGACGCAACGCGAAGAACCUUACCAAGGCUUGACAUACACCGGAAACGUCCAGAGAUGGGCGCCCCCUUGUGGUCGGUGUACAGGUGGUGCAUGGCUGUCGUCAGCUCGUGUCGUGAGAUGUUGGGUUAAGUCCCGCAACGAGCGCAACCCUUGUCCCGUGUUGCCAUCACGCCCUUGUGGUGCUGGGGACUCACGGGAGACCGCCGGGGUCAACUCGGA

**BB29** *Pseudomonas sp.*

AGCUUGAGCUCUUAGUGGCGCAGCUAACGCAUUAAGUUGACCGCCUGGGGAGUACGGCCGCAAGGUUAAAACUCAAAUGAAUUGACGGGGGCCCGCACAAGCGGUGGAGCAUGUGGUUUAAUUCGAAGCAACGCGAAGAACCUUACCAGGCCUUGACAUCCAAUGAACUUUCCAGAGAUGGAUUGGUGCCUUCGGGAACAUUGAGACAGGUGCUGCAUGGCUGUCGUCAGCUCGUGUCGUGAGAUGUUGGGUUAAGUCCCGUAACGAGCGCAACCCUUGUCCUUAGUUACCAGCACGUAAUGGUGGGCACUCUAAGGAGACUGCCGGUGACAAACCGGA

**BB30** *Pseudarthrobacter sp.*

GGUGUGGGGGACAUUCCACGUUUUCCGCGCCGUAGCUAACGCAUUAAGUGCCCCGCCUGGGGAGUACGGCCGCAAGGCUAAAACUCAAAGGAAUUGACGGGGGCCCGCACAAGCGGCGGAGCAUGCGGAUUAAUUCGAUGCAACGCGAAGAACCUUACCAAGGCUUGACAUGAACCGGAAAGACCUGGAAACAGGUGCCCCGCUUGCGGUCGGUUUACAGGUGGUGCAUGGUUGUCGUCAGCUCGUGUCGUGAGAUGUUGGGUUAAGUCCCGCAACGAGCGCAACCCUCGUUCUAUGUUGCCAGCACGUGAUGGUGGGGACUCAUAGGAGACUGCCGGGGUCAACUCGGA

**BB31** *Paenarthrobacter sp.*

UUCCACGUUUUUCCGCGCCGUAGCUUAACGCAUUUAAGUGCCCCGCCAGGGGAGUACGGCCGCAAGGCUAAAACUCAAAGGAAUUGACGGGGGCCCGCACAAGCGGCGGAGCAUGCGGAUUAAUUCGAUGCAACGCGAAGAACCUUACCAAGGCUUGACAUGAACCGGAAAGACCUGGAAACAGGUGCCCCGCUUGCGGUCGGUUUACAGGUGGUGCAUGGUUGUCGUCAGCUCGUGUCGUGAGAUGUUGGGUUAAGUCCCGCAACGAGCGCAACCCUCGUUCUAUGUUGCCAGCGGUUCGGCCGGGGACUCAUAGGAGACUGCCGGGGUCAACUCGGA

**BB32** *Rhizobium sp.*

GUCGGGCAGUAUACUGUUCGGUGGCGCAGCUAACGCAUUAAACAUUCCGCCUGGGGAGUACGGUCGCAAGAUUAAAACUCAAAGGAAUUGACGGGGGCCCGCACAAGCGGUGGAGCAUGUGGUUUAAUUCGAAGCAACGCGCAGAACCUUACCAGCCCUUGACAUGCCCGGCUACUUGCAGAGAUGCAAGGUUCCCUUCGGGGACCGGGACACAGGUGCUGCAUGGCUGUCGUCAGCUCGUGUCGUGAGAUGUUGGGUUAAGUCCCGCAACGAGCGCAACCCUCGCCCUUAGUUGCCAGCAUUCAGUUGGGCACUCUAAGGGGACUGCCGGUGAUAAGCCGAGA

**BB33** *Paenarthrobacter sp.*

GUGUGGGGGACAUUCCACGUUUUCCGCGCCGUAGCUAACGCAUUAAGUGCCCCGCCUGGGGAGUACGGCCGCAAGGCUAAAACUCAAAGGAAUUGACGGGGGCCCGCACAAGCGGCGGAGCAUGCGGAUUAAUUCGAUGCAACGCGAAGAACCUUACCAAGGCUUGACAUGAACCGGAAAGACCUGGAAACAGGUGCCCCGCUUGCGGUCGGUUUACAGGUGGUGCAUGGUUGUCGUCAGCUCGUGUCGUGAGAUGUUGGGUUAAGUCCCGCAACGAGCGCAACCCUCGUUCUAUGUUGCCAGCGGUUCGGCCGGGGACUCAUAGGAGACUGCCGGGGUCAACUCGGA

**BB34** *Pseudarthrobacter sp.*

AAGGUGUGGGGGAAUUCCACGUUUUCCGCGCCGUAGCUAACGCAUUAAGUCCCCCGCCNGGAGAGUGCCGCCGCAAGGCUAAAACUCAAAGGAAUUGACGGGGGCCCGCACAAGCGGCGGAGCAUGCGGAUUAAUUCGAUGCAACGCGAAGAACCUUACCAAGGCUUGACAUGAACCGGAAAGACCUGGAAACAGGUGCCCCGCUUGCGGUCGGUUUACAGGUGGUGCAUGGUUGUCGUCAGCUCGUGUCGUGAGAUGUUGGGUUAAGUCCCGCAACGAGCGCAACCCUCGUUCUAUGUUGCCAGCACGUGAUGGUGGGGACUCAUAGGAGACUGCCGGGGUCAACUCGGA

**BB35** *Paenarthrobacter sp.*

UGUGGGGGGAACAUUCCACGUUUUCCGCGCCGUAGCUAACGCAUUAAGUGCCCCGCCUGGGGAGUACGGCCGCAAGGCUAAAACUCAAAGGAAUUGACGGGGGCCCGCACAAGCGGCGGAGCAUGCGGAUUAAUUCGAUGCAACGCGAAGAACCUUACCAAGGCUUGACAUGAACCGGAAAGACCUGGAAACAGGUGCCCCGCUUGCGGUCGGUUUACAGGUGGUGCAUGGUUGUCGUCAGCUCGUGUCGUGAGAUGUUGGGUUAAGUCCCGCAACGAGCGCAACCCUCGUUCUAUGUUGCCAGCGGUUCGGCCGGGGACUCAUAGGAGACUGCCGGGGUCAACUCGGA

**BB36** *Rhizobium sp.*

UCGGGCAGUAUACUGUUCGGUGGCGCAGCUAACGCAUUAAACAUUCCGCCUGGGGAGUACGGUCGCAAGAUUAAAACUCAAAGGAAUUGACGGGGGCCCGCACAAGCGGUGGAGCAUGUGGUUUAAUUCGAAGCAACGCGCAGAACCUUACCAGCCCUUGACAUGCCCGGCUACUUGCAGAGAUGCAAGGUUCCCUUCGGGGACCGGGACACAGGUGCUGCAUGGCUGUCGUCAGCUCGUGUCGUGAGAUGUUGGGUUAAGUCCCGCAACGAGCGCAACCCUCGCCCUUAGUUGCCAGCAUUCAGUUGGGCACUCUAAGGGGACUGCCGGUGAUAAGCCGAGA

**BB37** *Paenarthrobacter sp.*

GGUGUGGGGGACAUUCCACGUUUUCCGCGCCGUAGCUAACGCAUUAAGUGCCCCGCCUGGGGAGUACGGCCGCAAGGCUAAAACUCAAAGGAAUUGACGGGGGCCCGCACAAGCGGCGGAGCAUGCGGAUUAAUUCGAUGCAACGCGAAGAACCUUACCAAGGCUUGACAUGAACCGGAAAGACCUGGAAACAGGUGCCCCGCUUGCGGUCGGUUUACAGGUGGUGCAUGGUUGUCGUCAGCUCGUGUCGUGAGAUGUUGGGUUAAGUCCCGCAACGAGCGCAACCCUCGUUCUAUGUUGCCAGCGGUUCGGCCGGGGACUCAUAGGAGACUGCCGGGGUCAACUCGGA

**BB38** *Nocardioides sp.*

AGCUAACGCAUUAAGCGCCCCGNNGNNNGUACGGCCGCAAGGCUAAAACUCAAAGGAAUUGACGGGGGCCCGCACAAGCGGCGGAGCAUGCGGAUUAAUUCGAUGCAACGCGAAGAACCUUACCUGGGUUUGACAUACACGGGAAGCCUGCAGAGAUGUGGGUCUCUUUGAUACUCGUGUACAGGUGGUGCAUGGCUGUCGUCAGCUCGUGUCGUGAGAUGUUGGGUUAAGUCCCGCAACGAGCGCAACCCUCGUUCCAUGUUGCCAGCGGGUUAUGCCGGGGACUCAUGGGAGACUGCCGGGGUCAACUCGGA

**BB39** *Streptomyces sp.*

GUGUUGGCGAACAUUCCACGUCGUCGGUGCCGCAGCUAACGCAUUAAGUUCCCCGCCUGGGGAGUACGGCCGCAAGGCUAAAACUCAAAGGAAUUGACGGGGGCCCGCACAAGCAGCGGAGCAUGUGGCUUAAUUCGACGCAACGCGAAGAACCUUACCAAGGCUUGACAUAUACCGGAAAGCAUCAGAGAUGGUGCCCCCCUUGUGGUCGGUAUACAGGUGGUGCAUGGCUGUCGUCAGCUCGUGUCGUGAGAUGUUGGGUUAAGUCCCGCAACGAGCGCAACCCUUGUUCUGUGUUGCCAGCAUGCCCUUCGGGGUGAUGGGGACUCACAGGAGACUGCCGGGGUCAACUCGGA

**BB40** *Pseudarthrobacter sp.*

GGUGUGGGGGACAUUCCACGUUUUCCGCGCCGUAGCUAACGCAUUAAGUGCCCCGCCNGUGNNAGUACGGCCGCAAGGCUAAAACUCAAAGGAAUUGACGGGGGCCCGCACAAGCGGCGGAGCAUGCGGAUUAAUUCGAUGCAACGCGAAGAACCUUACCAAGGCUUGACAUGAACCGGAAAGACCUGGAAACAGGUGCCCCGCUUGCGGUCGGUUUACAGGUGGUGCAUGGUUGUCGUCAGCUCGUGUCGUGAGAUGUUGGGUUAAGUCCCGCAACGAGCGCAACCCUCGUUCUAUGUUGCCAGCACGUGAUGGUGGGGACUCAUAGGAGACUGCCGGGGUCAACUCGGA

**BB41** *Paenarthrobacter sp.*

GGGGGAAUUCCACGGUUUUCCGCCGCCGUAGCUAACGCAUUAAGUGCCCCGCCUGGGGAGUACGGCCGCAAGGCUAAAACUCAAAGGAAUUGACGGGGGCCCGCACAAGCGGCGGAGCAUGCGGAUUAAUUCGAUGCAACGCGAAGAACCUUACCAAGGCUUGACAUGAACCGGAAAGACCUGGAAACAGGUGCCCCGCUUGCGGUCGGUUUACAGGUGGUGCAUGGUUGUCGUCAGCUCGUGUCGUGAGAUGUUGGGUUAAGUCCCGCAACGAGCGCAACCCUCGUUCUAUGUUGCCAGCGGUUCGGCCGGGGACUCAUAGGAGACUGCCGGGGUCAACUCGGA

**BB42** *Massilia sp.*

UGUUGGGUCUUAAUUGACUUAGUAACGCAGCUAACGCGUGAAGUAGACCGCCUGGCGAGUACGGUCGCAAGAUUAAAACUCAAAGGAAUUGACGGGGACCCGCACAAGCGGUGGAUGAUGUGGAUUAAUUCGAUGCAACGCGAAAAACCUUACCUACCCUUGACAUGGCAGGAAUCCCGGAGAGAUUUGGGAGUGCUCGAAAGAGAACCUGCACACAGGUGCUGCAUGGCUGUCGUCAGCUCGUGUCGUGAGAUGUUGGGUUAAGUCCCGCAACGAGCGCAACCCUUGUCAUUAGUUGCUACGCAAGAGCACUCUAAUGAGACUGCCGGUGACAAACCGGA

**BB43** *Mycobacterium sp.*

GGUGUGGGUUCCUUCCUUGGGAUCCGUGCCGUAGCUAACGCAUUAAGUACCCCGCCUGGGGAGUACGGCCGCAAGGCUAAAACUCAAAGGAAUUGACGGGGGCCCGCACAAGCGGCGGAGCAUGUGGAUUAAUUCGAUGCAACGCGAAGAACCUUACCUGGGUUUGACAUGCACAGGACGCUGGUAGAGAUAUCAGUUCCCUUGUGGCCUGUGUGCAGGUGGUGCAUGGCUGUCGUCAGCUCGUGUCGUGAGAUGUUGGGUUAAGUCCCGCAACGAGCGCAACCCCUAUCUUAUGUUGCCAGCGCGUUAUGGCGGGGACUCGUAAGAGACUGCCGGGGUCAACUCGGA

**BB45** *Phyllobacterium sp.*

UCGGGCAGUAUACUGUCUCGGUGGCGCAGCUAACGCAUUAAGCUCUCCGCCUGGGGAGUACGGUCGCAAGAUUAAAACUCAAAGGAAUUGACGGGGGCCCGCACAAGCGGUGGAGCAUGUGGUUUAAUUCGAAGCAACGCGCAGAACCUUACCAGCCCUUGACAUCCCGAUCGCGGUUACCAGAGAUGGUAUCCUUCAGUUAGGCUGGAUCGGUGACAGGUGCUGCAUGGCUGUCGUCAGCUCGUGUCGUGAGAUGUUGGGUUAAGUCCCGCAACGAGCGCAACCCUCGCCCUUAGUUGCCAUCAUUCAGUUGGGCACUCUAAGGGGACUGCCGGUGAUAAGCCGAGA

**Bacteria isolated from gramine treated soil (GB)**

**GB1** *Arthrobacter sp.*

GUGGGGGACAUUCCACGUUUUCCGCGCCGUAGCUAACGCAUUAAGUGCCCCGCCUGGGGAGUACGGCCGCAAGGCUAAAACUCAAAGGAAUUGACGGGGGCCCGCACAAGCGGCGGAGCAUGCGGAUUAAUUCGAUGCAACGCGAAGAACCUUACCAAGGCUUGACAUGAACCGGACCGCUGCAGAAAUGUGGUUUCCCCUUUGGGGCUGGUUUACAGGUGGUGCAUGGUUGUCGUCAGCUCGUGUCGUGAGAUGUUGGGUUAAGUCCCGCAACGAGCGCAACCCUCGUUCUAUGUUGCCAGCACGUGAUGGUGGGGACUCAUAGGAGACUGCCGGGGUCAACUCGGA

**GB2** *Pseudomonas sp.*

CUUAGUGGCGCAGCUAACGCAUUAAGUUGACCGCCUGGGGAGUACGGCCGCAAGGUUAAAACUCAAAUGAAUUGACGGGGGCCCGCACAAGCGGUGGAGCAUGUGGUUUAAUUCGAAGCAACGCGAAGAACCUUACCAGGCCUUGACAUCCAAUGAACUUUCCAGAGAUGGAUUGGUGCCUUCGGGAACAUUGAGACAGGUGCUGCAUGGCUGUCGUCAGCUCGUGUCGUGAGAUGUUGGGUUAAGUCCCGUAACGAGCGCAACCCUUGUCCUUAGUUACCAGCACGUUAUGGUGGGCACUCUAAGGAGACUGCCGGUGAC

**GB4** *Pseudomonas sp.*

CUUGAGCUCUUAGUGGCGCAGCUAACGCAUUAAGUUGACCGCCUGGGGAGUACGGCCGCAAGGUUAAAACUCAAAUGAAUUGACGGGGGCCCGCACAAGCGGUGGAGCAUGUGGUUUAAUUCGAAGCAACGCGAAGAACCUUACCAGGCCUUGACAUCCAAUGAACUUUCCAGAGAUGGAUUGGUGCCUUCGGGAACAUUGAGACAGGUGCUGCAUGGCUGUCGUCAGCUCGUGUCGUGAGAUGUUGGGUUAAGUCCCGUAACGAGCGCAACCCUUGUCCUUAGUUACCAGCACGUAAUGGUGGGCACUCUAAGGAGACUGC

**GB5** *Arthrobacter sp.*

GUGGGGGACAUUCCACGUUUUCCGCGCCGUAGCUAACGCAUUAAGUGCCCCGCCUGGGGAGUACGGCCGCAAGGCUAAAACUCAAAGGAAUUGACGGGGGCCCGCACAAGCGGCGGAGCAUGCGGAUUAAUUCGAUGCAACGCGAAGAACCUUACCAAGGCUUGACAUGAACCGGACCGCUGCAGAAAUGUGGUUUCCCCUUUGGGGCUGGUUUACAGGUGGUGCAUGGUUGUCGUCAGCUCGUGUCGUGAGAUGUUGGGUUAAGUCCCGCAACGAGCGCAACCCUCGUUCUAUGUUGCCAGCACGUGAUGGUGGGGACUCAUAGGAGACUGCCGGGGUCAACUCGGA

**GB6** *Pseudomonas sp.*

GCCGUUGGGAGCCUUGAGCUCUUAGUGGCGCAGCUAACGCAUUAAGUUGACCGCCUGGGGAGUACGGCCGCAAGGUUAAAACUCAAAUGAAUUGACGGGGGCCCGCACAAGCGGUGGAGCAUGUGGUUUAAUUCGAAGCAACGCGAAGAACCUUACCAGGCCUUGACAUCCAAUGAACUUUCCAGAGAUGGAUUGGUGCCUUCGGGAACAUUGAGACAGGUGCUGCAUGGCUGUCGUCAGCUCGUGUCGUGAGAUGUUGGGUUAAGUCCCGUAACGAGCGCAACCCUUGUCCUUAGUUACCAGCACGUAAUGGUGGGCACUCUAAGGAGACUGCCGGUGACAAACCGGA

**GB7** *Pseudarthrobacter sp.*

GGUGUGGGGGACAUUCCACGUUUUCCGCGCCGUAGCUAACGCAUUAAGUGCCCCGCCUGGGGAGUACGGCCGCAAGGCUAAAACUCAAAGGAAUUGACGGGGGCCCGCACAAGCGGCGGAGCAUGCGGAUUAAUUCGAUGCAACGCGAAGAACCUUACCAAGGCUUGACAUGAACCGGAAAGACCUGGAAACAGGUGCCCCGCUUGCGGUCGGUUUACAGGUGGUGCAUGGUUGUCGUCAGCUCGUGUCGUGAGAUGUUGGGUUAAGUCCCGCAACGAGCGCAACCCUCGUUCUAUGUUGCCAGCACGUGAUGGUGGGGACUCAUAGGAGACUGCCGGGGUCAACUCGGA

**GB8** *Pseudomonas sp.*

CGUUGGGAGCCUUGAGCUCUUAGUGGCGCAGCUAACGCAUUAAGUUGACCGCCUGGGGAGUACGGCCGCAAGGUUAAAACUCAAAUGAAUUGACGGGGGCCCGCACAAGCGGUGGAGCAUGUGGUUUAAUUCGAAGCAACGCGAAGAACCUUACCAGGCCUUGACAUCCAAUGAACUUUCCAGAGAUGGAUGGGUGCCUUCGGGAACAUUGAGACAGGUGCUGCAUGGCUGUCGUCAGCUCGUGUCGUGAGAUGUUGGGUUAAGUCCCGUAACGAGCGCAACCCUUGUCCUUAGUUACCAGCACGUAAUGGUGGGCACUCUAAGGAGACUGCCGGUGACAAACCGG

**GB9** *Arthrobacter sp.*

GUGUGGGGGGACAUUCCACGUUUUCCGCGCCGUAGCUAACGCAUUAAGUGCCCCGCCUGGGGAGUACGGCCGCAAGGCUAAAACUCAAAGGAAUUGACGGGGGCCCGCACAAGCGGCGGAGCAUGCGGAUUAAUUCGAUGCAACGCGAAGAACCUUACCAAGGCUUGACAUGAACCGGACCGCUGCAGAAAUGUGGUUUCCCCUUUGGGGCUGGUUUACAGGUGGUGCAUGGUUGUCGUCAGCUCGUGUCGUGAGAUGUUGGGUUAAGUCCCGCAACGAGCGCAACCCUCGUUCUAUGUUGCCAGCACGUGAUGGUGGGGACUCAUAGGAGACUGCCGGGGUCAACUCGGA

**GB10** *Pseudomonas sp.*

GCCGUUGGGAGCCUUGAGCUCUUAGUGGCGCAGCUAACGCAUUAAGUUGACCGCCUGGGGAGUACGGCCGCAAGGUUAAAACUCAAAUGAAUUGACGGGGGCCCGCACAAGCGGUGGAGCAUGUGGUUUAAUUCGAAGCAACGCGAAGAACCUUACCAGGCCUUGACAUCCAAUGAACUUUCCAGAGAUGGAUUGGUGCCUUCGGGAACAUUGAGACAGGUGCUGCAUGGCUGUCGUCAGCUCGUGUCGUGAGAUGUUGGGUUAAGUCCCGUAACGAGCGCAACCCUUGUCCUUAGUUACCAGCACGUAAUGGUGGGCACUCUAAGGAGACUGCCGGUGACAAACCGGA

**GB11** *Pseudarthrobacter sp.*

GUGUGGGGGGACAUUCCACGUUUUCCGCGCCGUAGCUAACGCAUUAAGUGCCCCGCCUGGGGAGUACGGCCGCAAGGCUAAAACUCAAAGGAAUUGACGGGGGCCCGCACAAGCGGCGGAGCAUGCGGAUUAAUUCGAUGCAACGCGAAGAACCUUACCAAGGCUUGACAUGAACCGGUAACGCCUGGAAACAGGUGCCCCGCUUGCGGUCGGUUUACAGGUGGUGCAUGGUUGUCGUCAGCUCGUGUCGUGAGAUGUUGGGUUAAGUCCCGCAACGAGCGCAACCCUCGUUCUAUGUUGCCAGCACGUGAUGGUGGGGACUCAUAGGAGACUGCCGGGGUCAACUCGGA

**GB12** *Pseudomonas sp.*

UUGGGAGCCUUGAGCUCUUAGUGGCGCAGCUAACGCAUUAAGUUGACCGCCUGGGGAGUACGGCCGCAAGGUUAAAACUCAAAUGAAUUGACGGGGGCCCGCACAAGCGGUGGAGCAUGUGGUUUAAUUCGAAGCAACGCGAAGAACCUUACCAGGCCUUGACAUCCAAUGAACUUUCCAGAGAUGGAUGGGUGCCUUCGGGAACAUUGAGACAGGUGCUGCAUGGCUGUCGUCAGCUCGUGUCGUGAGAUGUUGGGUUAAGUCCCGUAACGAGCGCAACCCUUGUCCUUAGUUACCAGCACGUAAUGGUGGGCACUCUAAGGAGACUGCCGGUGACAAACCGGA

**GB13** *Pseudomonas sp.*

AGCCGUUGGGAGCCUUGAGCUCUUAGUGGCGCAGCUAACGCAUUAAGUUGACCGCCUGGGGAGUACGGCCGCAAGGUUAAAACUCAAAUGAAUUGACGGGGGCCCGCACAAGCGGUGGAGCAUGUGGUUUAAUUCGAAGCAACGCGAAGAACCUUACCAGGCCUUGACAUCCAAUGAACUUUCCAGAGAUGGAUUGGUGCCUUCGGGAACAUUGAGACAGGUGCUGCAUGGCUGUCGUCAGCUCGUGUCGUGAGAUGUUGGGUUAAGUCCCGUAACGAGCGCAACCCUUGUCCUUAGUUACCAGCACGUAAUGGUGGGCACUCUAAGGAGACUGCCGGUGACAAACCGGA

**GB14** *Paenarthrobacter sp.*

GUGUGGGGGACAUUCCACGUUUUCCGCGCCGUAGCUAACGCAUUAAGUGCCCCGCCUGGGGAGUACGGCCGCAAGGCUAAAACUCAAAGGAAUUGACGGGGGCCCGCACAAGCGGCGGAGCAUGCGGAUUAAUUCGAUGCAACGCGAAGAACCUUACCAAGGCUUGACAUGAACCGGAAAGACCUGGAAACAGGUGCCCCGCUUGCGGUCGGUUUACAGGUGGUGCAUGGUUGUCGUCAGCUCGUGUCGUGAGAUGUUGGGUUAAGUCCCGCAACGAGCGCAACCCUCGUUCUAUGUUGCCAGCGGUUCGGCCGGGGACUCAUAGGAGACUGCCGGGGUCAACUCAGA

**GB15** *Pseudomonas sp.*

CCGUUGGGAGCCUUGAGCUCUUAGUGGCGCAGCUAACGCAUUAAGUUGACCGCCUGGGGAGUACGGCCGCAAGGUUAAAACUCAAAUGAAUUGACGGGGGCCCGCACAAGCGGUGGAGCAUGUGGUUUAAUUCGAAGCAACGCGAAGAACCUUACCAGGCCUUGACAUCCAAUGAACUUUCCAGAGAUGGAUUGGUGCCUUCGGGAACAUUGAGACAGGUGCUGCAUGGCUGUCGUCAGCUCGUGUCGUGAGAUGUUGGGUUAAGUCCCGUAACGAGCGCAACCCUUGUCCUUAGUUACCAGCACGUAAUGGUGGGCACUCUAAGGAGACUGCCGGUGACAAACCGGA

**GB16** *Streptomyces sp.*

AGGUGUUGGCGACAUUCCACGUCGUCGGUGCCGCAGCUAACGCAUUAAGUUCCCCGCCUGGGGAGUACGGCCGCAAGGCUAAAACUCAAAGGAAUUGACGGGGGCCCGCACAAGCAGCGGAGCAUGUGGCUUAAUUCGACGCAACGCGAAGAACCUUACCAAGGCUUGACAUCGCCCGGAAAGCAUCAGAGAUGGUGCCCCCCUUGUGGUCGGGUGACAGGUGGUGCAUGGCUGUCGUCAGCUCGUGUCGUGAGAUGUUGGGUUAAGUCCCGCAACGAGCGCAACCCUUGUUCUGUGUUGCCAGCAUGCCCUUCGGGGUGAUGGGGACUCACAGGAGACUGCCGGGGU

**GB17** *Arthrobacter sp.*

UUCCACGUUUUCCGCGCCGUAGCUAACGCAUUAAGUGCCCCGCCUGGGGAGUACGGCCGCAAGGCUAAAACUCAAAGGAAUUGACGGGGGCCCGCACAAGCGGCGGAGCAUGCGGAUUAAUUCGAUGCAACGCGAAGAACCUUACCAAGGCUUGACAUGGACCGGACCGCCGCAGAAAUGUGGUUUCCCCUUUGGGGCCGGUUCACAGGUGGUGCAUGGUUGUCGUCAGCUCGUGUCGUGAGAUGUUGGGUUAAGUCCCGCAACGAGCGCAACCCUCGUUCCAUGUUGCCAGCGCGUAAUGGCGGGGACUCAUGGGAGACUGCCGGGGUCAACUCGGA

**GB18** *Arthobacter sp.*

GGUGUGGGGGACAUUCCACGUUUUCCGCGCCGUAGCUAACGCAUUAAGUGCCCCGCCUGGGGAGUACGGCCGCAAGGCUAAAACUCAAAGGAAUUGACGGGGGCCCGCACAAGCGGCGGAGCAUGCGGAUUAAUUCGAUGCAACGCGAAGAACCUUACCAAGGCUUGACAUGGACCGGACCGCCGCAGAAAUGUGGUUUCCCCUUUGGGGCCGGUUCACAGGUGGUGCAUGGUUGUCGUCAGCUCGUGUCGUGAGAUGUUGGGUUAAGUCCCGCAACGAGCGCAACCCUCGUUCCAUGUUGCCAGCGCGUAAUGGCGGGGACUCAUGGGAGACUGCCGGGG

**GB19** *Streptomyces sp.*

UCCACGUCGUCGGUGCCGCAGCUAACGCAUUAAGUUCCCCGCCUGGGGAGUACGGCCGCAAGGCUAAAACUCAAAGGAAUUGACGGGGGCCCGCACAAGCAGCGGAGCAUGUGGCUUAAUUCGACGCAACGCGAAGAACCUUACCAAGGCUUGACAUAUACCGGAAAGCAUCAGAGAUGGUGCCCCCCUUGUGGUCGGUAUACAGGUGGUGCAUGGCUGUCGUCAGCUCGUGUCGUGAGAUGUUGGGUUAAGUCCCGCAACGAGCGCAACCCUUGUUCUGUGUUGCCAGCAUGCCCUUCGGGGUGAUGGGGACUCACAGGAGACU

**GB20** *Arthrobacter sp.*

UCCACGUUUUCCGCGCCGUAGCUAACGCAUUAAGUGCCCCGCCUGGGGAGUACGGCCGCAAGGCUAAAACUCAAAGGAAUUGACGGGGGCCCGCACAAGCGGCGGAGCAUGCGGAUUAAUUCGAUGCAACGCGAAGAACCUUACCAAGGCUUGACAUGGACCGGACCGCCGCAGAAAUGUGGUUUCCCCUUUGGGGCCGGUUCACAGGUGGUGCAUGGUUGUCGUCAGCUCGUGUCGUGAGAUGUUGGGUUAAGUCCCGCAACGAGCGCAACCCUCGUUCCAUGUUGCCAGCGCGUAAUGGCGGGGACUCAUGGGAGACUGCCGGG

**GB21** *Arthrobacter sp.*

UUCCACGUUUUCCGCGCCGUAGCUAACGCAUUAAGUGCCCCGCCUGGGGAGUACGGCCGCAAGGCUAAAACUCAAAGGAAUUGACGGGGGCCCGCACAAGCGGCGGAGCAUGCGGAUUAAUUCGAUGCAACGCGAAGAACCUUACCAAGGCUUGACAUGGACCGGACCGCCGCAGAAAUGUGGUUUCCCCUUUGGGGCCGGUUCACAGGUGGUGCAUGGUUGUCGUCAGCUCGUGUCGUGAGAUGUUGGGUUAAGUCCCGCAACGAGCGCAACCCUCGUUCCAUGUUGCCAGCGCGUAAUGGCGGGGACUCAUGGGAGACUGCCGGGG

**Bacteria isolated from quercetin treated soil (QB)**

**QB1** *Arthrobacter sp.*

CCCUGGUAGUCCAUGCCGUAAACGUUGGGCACUAGGUGUGGGGGACAUUCCACGUUUUCCGCGCCGUAGCUAACGCAUUAAGUGCCCCGCCUGGGGAGUACGGCCGCAAGGCUAAAACUCAAAGGAAUUGACGGGGGCCCGCACAAGCGGCGGAGCAUGCGGAUUAAUUCGAUGCAACGCGAAGAACCUUACCAAGGCUUGACAUGAACCGGACCGCUGCAGAAAUGUGGUUUCCCCUUUGGGGCUGGUUUACAGGUGGUGCAUGGUUGUCGUCAGCUCGUGUCGUGAGAUGUUGGGUUAAGUCCCGCAACGAGCGCAACCCUCGUUCUAUGUUGCCAGCACGUGAUGGU

**QB2** *Pseudarthrobacter sp.*

UAGUCCAUGCCGUAAACGUUGGGCACUAGGUGUGGGGGACAUUCCACGUUUUCCGCGCCGUAGCUAACGCAUUAAGUGCCCCGCCUGGGGAGUACGGCCGCAAGGCUAAAACUCAAAGGAAUUGACGGGGGCCCGCACAAGCGGCGGAGCAUGCGGAUUAAUUCGAUGCAACGCGAAGAACCUUACCAAGGCUUGACAUGAACCGGAAAGACCUGGAAACAGGUCCCCCACUUGUGGUCGGUUUACAGGUGGUGCAUGGUUGUCGUCAGCUCGUGUCGUGAGAUGUUGGGUUAAGUCCCGCAACGAGCGCAACCCUCGUUCUAUGUUGCCAGCACGUGAUGGU

**QB3** *Pseudarthrobacter sp.*

CCCUGGUAGUCCAUGCCGUAAACGUUGGGCACUAGGUGUGGGGGACAUUCCACGUUUUCCGCGCCGUAGCUAACGCAUUAAGUGCCCCGCCUGGGGAGUACGGCCGCAAGGCUAAAACUCAAAGGAAUUGACGGGGGCCCGCACAAGCGGCGGAGCAUGCGGAUUAAUUCGAUGCAACGCGAAGAACCUUACCAAGGCUUGACAUGAACCGGAAAGACCUGGAAACAGGUCCCCCACUUGUGGUCGGUUUACAGGUGGUGCAUGGUUGUCGUCAGCUCGUGUCGUGAGAUGUUGGGUUAAGUCCCGCAACGAGCGCAACCCUCGUUCUAUGUUGCCAGCACGUGAUGGU

**QB4** *Pseudarthrobacter sp.*

CCCUGGUAGUCCAUGCCGUAAACGUUGGGCACUAGGUGUGGGGGACAUUCCACGUUUUCCGCGCCGUAGCUAACGCAUUAAGUGCCCCGCCUGGGGAGUACGGCCGCAAGGCUAAAACUCAAAGGAAUUGACGGGGGCCCGCACAAGCGGCGGAGCAUGCGGAUUAAUUCGAUGCAACGCGAAGAACCUUACCAAGGCUUGACAUGAACCGGAAAGACCUGGAAACAGGUCCCCCACUUGUGGUCGGUUUACAGGUGGUGCAUGGUUGUCGUCAGCUCGUGUCGUGAGAUGUUGGGUUAAGUCCCGCAACGAGCGCAACCCUCGUUCUAUGUUGCCAGCACGUGAUGGUGGGAC

**QB5** *Pseudarthrobacter sp.*

CCCUGGUAGUCCAUGCCGUAAACGUUGGGCACUAGGUGUGGGGGACAUUCCACGUUUUCCGCGCCGUAGCUAACGCAUUAAGUGCCCCGCCUGGGGAGUACGGCCGCAAGGCUAAAACUCAAAGGAAUUGACGGGGGCCCGCACAAGCGGCGGAGCAUGCGGAUUAAUUCGAUGCAACGCGAAGAACCUUACCAAGGCUUGACAUGAACCGGAAAGACCUGGAAACAGGUCCCCCACUUGUGGUCGGUUUACAGGUGGUGCAUGGUUGUCGUCAGCUCGUGUCGUGAGAUGUUGGGUUAAGUCCCGCAACGAGCGCAACCCUCGUUCUAUGUUGCCAGCACGUGAUGG

**QB6** *Pseudarthrobacter sp.*

CCCUGGUAGUCCAUGCCGUAAACGUUGGGCACUAGGUGUGGGGGACAUUCCACGUUUUCCGCGCCGUAGCUAACGCAUUAAGUGCCCCGCCUGGGGAGUACGGCCGCAAGGCUAAAACUCAAAGGAAUUGACGGGGGCCCGCACAAGCGGCGGAGCAUGCGGAUUAAUUCGAUGCAACGCGAAGAACCUUACCAAGGCUUGACAUGAACCGGAAAGACCUGGAAACAGGUCCCCCACUUGUGGUCGGUUUACAGGUGGUGCAUGGUUGUCGUCAGCUCGUGUCGUGAGAUGUUGGGUUAAGUCCCGCAACGAGCGCAACCCUCGUUCUAUGUUGCCAGCACGUGAUGGUGGG

**QB7** *Pseudarthrobacter sp.*

CCCUGGUAGUCCAUGCCGUAAACGUUGGGCACUAGGUGUGGGGGACAUUCCACGUUUUCCGCGCCGUAGCUAACGCAUUAAGUGCCCCGCCUGGGGAGUACGGCCGCAAGGCUAAAACUCAAAGGAAUUGACGGGGGCCCGCACAAGCGGCGGAGCAUGCGGAUUAAUUCGAUGCAACGCGAAGAACCUUACCAAGGCUUGACAUGAACCGGAAAGACCUGGAAACAGGUCCCCCACUUGUGGUCGGUUUACAGGUGGUGCAUGGUUGUCGUCAGCUCGUGUCGUGAGAUGUUGGGUUAAGUCCCGCAACGAGCGCAACCCUCGUUCUAUGUUGCCAGCACGUGAUGGUGGG

**QB8** *Novosphingobium sp.*

CCUGGUAGUCCACGCCGUAAACGAUGAUAACUAGCUGUCCGGGUACUUGGUACUUGGGUGGCGCAGCUAACGCAUUAAGUUAUCCGCCUGGGGAGUACGGUCGCAAGAUUAAAACUCAAAGGAAUUGACGGGGGCCUGCACAAGCGGUGGAGCAUGUGGUUUAAUUCGAAGCAACGCGCAGAACCUUACCAGCGUUUGACAUGCCGGUCGCGGAUUUGGGAGACCAUUUCCUUCAGUUCGGCUGGACCGUGCACAGGUGCUGCAUGGCUGUCGUCAGCUCGUGUCGUGAGAUGUUGGGUUAAGUCCCGCAACGAGCGCAACCCUCGUCCUUAGUUGCCAGCAUU

**QB10** *Nocardioides sp.*

GUAGUCCACACCGUAAACGUUGGGCGCUAGGUGUGGGGCUCAUUCCACGAGUUCCGUGCCGCAGCUAACGCAUUAAGCGCCCCGCCUGGGGAGUACGGCCGCAAGGCUAAAACUCAAAGGAAUUGACGGGGGCCCGCACAAGCGGCGGAGCAUGCGGAUUAAUUCGAUGCAACGCGAAGAACCUUACCUGGGUUUGACAUAUGCCGGAAAGCAUCAGAGAUGGUGCCCCUUUUUGUCGGUAUACAGGUGGUGCAUGGCUGUCGUCAGCUCGUGUCGUGAGAUGUUGGGUUAAGUCCCGCAACGAGCGCAACCCUCGUCUUAUGUUGCCAGCACGUAAUGGUGGGA

**QB11** *Novosphingobium sp.*

CCCUGGUAGUCCACGCCGUAAACGAUGAUAACUAGCUGUCCGGGUACUUGGUACUUGGGUGGCGCAGCUAACGCAUUAAGUUAUCCGCCUGGGGAGUACGGUCGCAAGAUUAAAACUCAAAGGAAUUGACGGGGGCCUGCACAAGCGGUGGAGCNUGUGGUUUAAUUCGAAGCAACGCGCAGAACCUUACCAGCGUUUGACAUGCCGGUCGCGGAUUUGGGAGACCAUUUCCUUCAGUUCGGCUGGACCGUGCACAGGUGCUGCAUGGCUGUCGUCAGCUCGUGUCGUGAGAUGUUGGGUUAAGUCCCGCAACGAGCGCAACCCUCGUCCUUAGUUGCCAGCAUUG

**QB12** *Pseudarthrobacter sp.*

CCCUGGUAGUCCAUGCCGUAAACGUUGGGCACUAGGUGUGGGGGACAUUCCACGUUUUCCGCGCCGUAGCUAACGCAUUAAGUGCCCCGCCUGGGGAGUACGGCCGCAAGGCUAAAACUCAAAGGAAUUGACGGGGGCCCGCACAAGCGGCGGAGCAUGCGGAUUAAUUCGAUGCAACGCGAAGAACCUUACCAAGGCUUGACAUGAACCGGAAAGACCUGGAAACAGGUCCCCCACUUGUGGUCGGUUUACAGGUGGUGCAUGGUUGUCGUCAGCUCGUGUCGUGAGAUGUUGGGUUAAGUCCCGCAACGAGCGCAACCCUCGUUCUAUGUUGCCAGCACGUGAUGG

**QB13** *Pseudarthrobacter sp.*

GGUAGUCCAUGCCGUAAACGUUGGGCACUAGGUGUGGGGGACAUUCCACGUUUUCCGCGCCGUAGCUAACGCAUUAAGUGCCCCGCCUGGGGAGUACGGCCGCAAGGCUAAAACUCAAAGGAAUUGACGGGGGCCCGCACAAGCGGCGGAGCAUGCGGAUUAAUUCGAUGCAACGCGAAGAACCUUACCAAGGCUUGACAUGAACCGGAAAGACCUGGAAACAGGUCCCCCACUUGUGGUCGGUUUACAGGUGGUGCAUGGUUGUCGUCAGCUCGUGUCGUGAGAUGUUGGGUUAAGUCCCGCAACGAGCGCAACCCUCGUUCUAUGUUGCCAGCACGUGAUGG

**QB14** *Pseudarthrobacter sp.*

CCUGGUAGUCCAUGCCGUAAACGUUGGGCACUAGGUGUGGGGGACAUUCCACGUUUUCCGCGCCGUAGCUAACGCAUUAAGUGCCCCGCCUGGGGAGUACGGCCGCAAGGCUAAAACUCAAAGGAAUUGACGGGGGCCCGCACAAGCGGCGGAGCAUGCGGAUUAAUUCGAUGCAACGCGAAGAACCUUACCAAGGCUUGACAUGGACUGGAAAUACCUGGAAACAGGUGCCCCGCUUGCGGCCGGUUCACAGGUGGUGCAUGGUUGUCGUCAGCUCGUGUCGUGAGAUGUUGGGUUAAGUCCCGCAACGAGCGCAACCCUCGUUCUAUGUUGCCAGCGCGUUAUGGCG

**QB15** *Novosphingobium sp.*

CCUGGUAGUCCACGCCGUAAACGAUGAUAACUAGCUGUCCGGGUACUUGGUACUUGGGUGGCGCAGCUAACGCAUUAAGUUAUCCGCCUGGGGAGUACGGUCGCAAGAUUAAAACUCAAAGGAAUUGACGGGGGCCUGCACAAGCGGUGGAGCAUGUGGUUUAAUUCGAAGCAACGCGCAGAACCUUACCAGCGUUUGACAUGCCGGUCGCGGAUUUGGGAGACCAUUUCCUUCAGUUCGGCUGGACCGUGCACAGGUGCUGCAUGGCUGUCGUCAGCUCGUGUCGUGAGAUGUUGGGUUAAGUCCCGCAACGAGCGCAACCCUCGUCCUUAGUUGCCAGCAUUUA

**QB16** *Pseudomonas sp.*

UGGUAGUCCACGCCGUAAACGAUGUCGACUAGCCGUUGGGGUCCUUGAGACUUUAGUGGCGCAGCUAACGCAUUAAGUCGACCGCCUGGGGAGUACGGCCGCAAGGUUAAAACUCAAAUGAAUUGACGGGGGCCCGCACAAGCGGUGGAGCAUGUGGUUUAAUUCGAAGCAACGCGAAGAACCUUACCUGGCCUUGACAUGCUGAGAACUUUCUAGAGAUAGAUUGGUGCCUUCGGGAACUCAGACACAGGUGCUGCAUGGCUGUCGUCAGCUCGUGUCGUGAGAUGUUGGGUUAAGUCCCGUAACGAGCGCAACCCUUGUCCUUAGUUACCAGCACGUUAUGG

**QB17** *Arthrobacter sp.*

UGGUAGUCCAUGCCGUAAACGUUGGGCACUAGGUGUGGGGGACAUUCCACGUUUUCCGCGCCGUAGCUAACGCAUUAAGUGCCCCGCCUGGGGAGUACGGCCGCAAGGCUAAAACUCAAAGGAAUUGACGGGGGCCCGCACAAGCGGCGGAGCAUGCGGAUUAAUUCGAUGCAACGCGAAGAACCUUACCAAGGCUUGACAUGAACCGGACCGCUGCAGAAAUGUGGUUUCCCCUUUGGGGCUGGUUUACAGGUGGUGCAUGGUUGUCGUCAGCUCGUGUCGUGAGAUGUUGGGUUAAGUCCCGCAACGAGCGCAACCCUCGUUCUAUGUUGCCAGCACGUGAUGGU

**QB18** *Pseudarthrobacter sp.*

CCCUGGUAGUCCAUGCCGUAAACGUUGGGCACUAGGUGUGGGGGACAUUCCACGUUUUCCGCGCCGUAGCUAACGCAUUAAGUGCCCCGCCUGGGGAGUACGGCCGCAAGGCUAAAACUCAAAGGAAUUGACGGGGGCCCGCACAAGCGGCGGAGCAUGCGGAUUAAUUCGAUGCAACGCGAAGAACCUUACCAAGGCUUGACAUGGACUGGAAAUACCUGGAAACAGGUGCCCCGCUUGCGGCCGGUUCACAGGUGGUGCAUGGUUGUCGUCAGCUCGUGUCGUGAGAUGUUGGGUUAAGUCCCGCAACGAGCGCAACCCUCGUUCUAUGUUGCCAGCGCGUGAUGGCGGG

**QB19** *Novosphingobium sp.*

CCCUGGUAGUCCACGCCGUAAACGAUGAUAACUAGCUGUCCGGGUACUUGGUACUUGGGUGGCGCAGCUAACGCAUUAAGUUAUCCGCCUGGGGAGUACGGUCGCAAGAUUAAAACUCAAAGGAAUUGACGGGGGCCUGCACAAGCGGUGGAGCAUGUGGUUUAAUUCGAAGCAACGCGCAGAACCUUACCAGCGUUUGACAUGCCGGUCGCGGAUUUGGGAGACCAUUUCCUUCAGUUCGGCUGGACCGUGCACAGGUGCUGCAUGGCUGUCGUCAGCUCGUGUCGUGAGAUGUUGGGUUAAGUCCCGCAACGAGCGCAACCCUCGUCCUUAGUUGCCAGCAUU

**QB20** *Novosphingobium sp.*

GGUACUUGGUACUUGGGUGGCGCAGCUAACGCAUUAAGUUAUCCGCCUGGGGAGUACGGUCGCAAGAUUAAAACUCAAAGGAAUUGACGGGGGCCUGCACAAGCGGUGGAGCAUGUGGUUUAAUUCGAAGCAACGCGCAGAACCUUACCAGCGUUUGACAUGCCGGUCGCGGAUUUGGGAGACCAUUUCCUUCAGUUCGGCUGGACCGUGCACAGGUGCUGCAUGGCUGUCGUCAGCUCGUGUCGUGAGAUGUUGGGUUAAGUCCCGCAACGAGCGCAACCCUCGUCCUUAGUUGCCAGCAUUGAGUUGGGCACUCUAAGGAAACUGCCGGUGAUAAGCCGGAGG

**QB21** *Pseudomonas sp.*

AAGCCGUUGGGAGCCUUGAGCUCUUAGUGGCGCAGCUAACGCAUUAAGUUGACCGCCUGGGGAGUACGGCCGCAAGGUUAAAACUCAAAUGAAUUGACGGGGGCCCGCACAAGCGGUGGAGCAUGUGGUUUAAUUCGAAGCAACGCGAAGAACCUUACCAGGCCUUGACAUCCAAUGAACUUUCCAGAGAUGGAUUGGUGCCUUCGGGAACAUUGAGACAGGUGCUGCAUGGCUGUCGUCAGCUCGUGUCGUGAGAUGUUGGGUUAAGUCCCGUAACGAGCGCAACCCUUGUCCUUAGUUACCAGCACGUUAUGGUGGGCACUCUAAGGAUACUGCCGGU

**QB22** *Arthrobacter sp.*

UCCACGUUUUCCGCGCCGUAGCUAACGCAUUAAGUGCCCCGCCUGGGGAGUACGGCCGCAAGGCUAAAACUCAAAGGAAUUGACGGGGGCCCGCACAAGCGGCGGAGCAUGCGGAUUAAUUCGAUGCAACGCGAAGAACCUUACCAAGGCUUGACAUGGACCGGACCGCCGCAGAAAUGUGGUUUCCCCUUUGGGGCCGGUUCACAGGUGGUGCAUGGUUGUCGUCAGCUCGUGUCGUGAGAUGUUGGGUUAAGUCCCGCAACGAGCGCAACCCUCGUUCCAUGUUGCCAGCGCGUAAUGGCGGGGACUCAUGGGAGACUGCCGGGG

**QB23** *Arthrobacter sp.*

CACGUUUUCCGCGCCGUAGCUAACGCAUUAAGUGCCCCGCCUGGGGAGUACGGCCGCAAGGCUAAAACUCAAAGGAAUUGACGGGGGCCCGCACAAGCGGCGGAGCAUGCGGAUUAAUUCGAUGCAACGCGAAGAACCUUACCAAGGCUUGACAUGGACCGGACCGCCGCAGAAAUGUGGUUUCCCCUUUGGGGCCGGUUCACAGGUGGUGCAUGGUUGUCGUCAGCUCGUGUCGUGAGAUGUUGGGUUAAGUCCCGCAACGAGCGCAACCCUCGUUCCAUGUUGCCAGCGCGUAAUGGCGGGGACUCAUGGGAGACUGCCGGGG
