## Supplementary material for "Differential impact of plant secondary metabolites on the soil microbiota": Figure S1

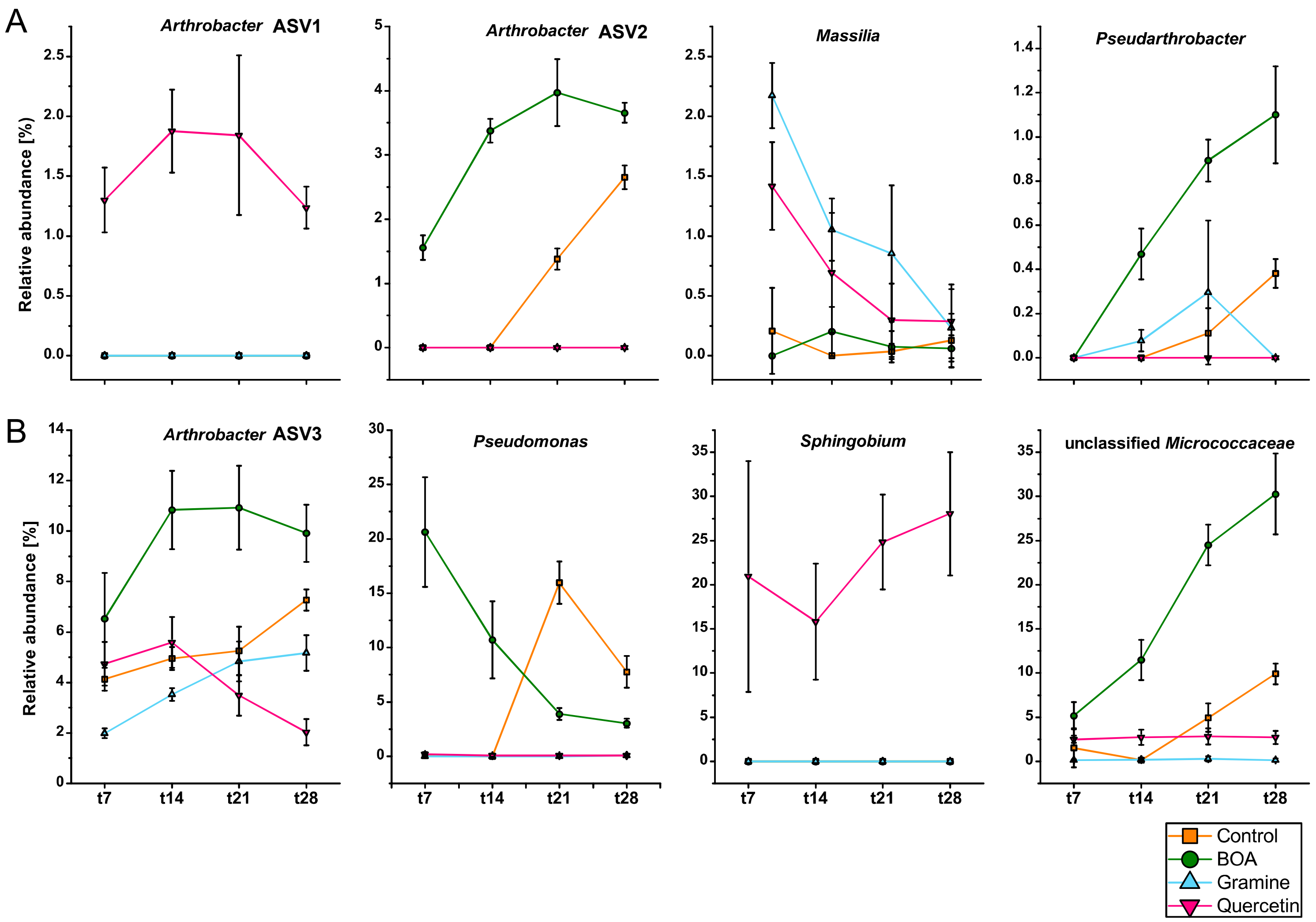


**Figure S1. Alterations in relative abundances of individual ASVs during treatment with plant metabolites.**

The changes in relative abundances of representative ASVs based on amplicon sequencing are shown for the different days of treatments with BOA, gramine or quercetin.

**A,** Representative ASVs showing typical dynamics of responses to the different plant metabolites.

**B,** Four ASVs that were highly abundant at least during one soil treatment.
